# Modulating the left inferior frontal cortex by task domain, cognitive challenge and tDCS

**DOI:** 10.1101/2021.02.05.429968

**Authors:** Davide Nardo, Katerina Pappa, John Duncan, Peter Zeidman, Martina F. Callaghan, Alexander Leff, Jennifer Crinion

## Abstract

The left inferior frontal cortex (LIFC) is a key region for spoken language processing, but its neurocognitive architecture remains controversial. Here we assess the domain-generality vs. domain-specificity of the LIFC from behavioural, functional neuroimaging and neuromodulation data. Using concurrent fMRI and transcranial direct current stimulation (tDCS) delivered to the LIFC, we investigated how brain activity and behavioural performance are modulated by task domain (naming vs. non-naming), cognitive challenge (low vs. high), and tDCS (anodal vs. sham). The data revealed: (1) co-existence of neural signatures both common and distinct across tasks within the LIFC; (2) domain-preferential effects of task (naming); (3) significant tDCS modulations of activity in a LIFC sub-region selectively during high-challenge naming. The presence of both domain-specific and domain-general signals, and the existence of a gradient of activation where naming relied more on sub-regions within the LIFC, may help reconcile both perspectives on spoken language processing.

## INTRODUCTION

Since Paul Broca’s seminal discovery of the localisation of expressive aphasia in the damaged brain more than 150 years ago, the left inferior frontal cortex (LIFC) has been considered a key brain region for speech function. In the last three decades, the advent of functional imaging has provided plenty of evidence supporting the relationship between speech production and activity in the LIFC in healthy subjects (e.g., for a review see Price, 2012), also showing that the LIFC is implicated in other key aspects of language processing, such as comprehension, syntax, and semantics (Dapretto & Bookheimer, 1999; Noppeney et al., 2004; Tyler et al., 2011; Rodd et al., 2015).

However, critically, functional imaging research has shown that regions within the LIFC also contribute to an executive function network activated by many non-linguistic, cognitively challenging tasks (e.g., see Bartley et al., 2018; Camilleri et al., 2018). Whether language, as a mental process is domain-general (i.e., shares a single underlying resource across many cognitive functions or tasks) or domain-specific (i.e., relies on independent components) is a broad question (cf. Petkov & Marslen-Wilson, 2018) that is pertinent to many areas of psychology. In cognitive neuroscience, whether the LIFC might be part of a network supporting domain-general (i.e., multiple cognitively challenging tasks), rather than domain-specific (i.e., mainly linguistic-related tasks), is hotly debated (Duncan, 2010; Fedorenko et al., 2012; Fedorenko & Thompson-Schill, 2014; Geranmayeh et al., 2014). In this paper, we focus on the issue of domain-specificity vs. domain-generality of neurocognitive substrates supporting spoken language in the LIFC.

Initial evidence of a domain-general role for the LIFC comes from a series of functional imaging studies investigating the issue of specificity vs. generality within the language domain, i.e. using verbal stimulus material. These studies tried to disentangle whether the LIFC (or any sub-region within it) is associated with specific aspects of linguistic processing (e.g., phonology, syntax, semantics), or rather if its activity is dynamically associated with cognitive demand. For instance, activity in the LIFC may not be associated with semantic retrieval *per se*, but rather with general cognitive selection demands, such as when faced with many competing alternative responses (e.g., when naming a picture of a dog you choose to say either /animal/, /dog/, /pet/, /Dalmatian/, /Fido/etc.; cf. Thompson-Schill et al., 1997). Other studies suggest that activity in this region may be related to increased cognitive effort due to conflict and/or ambiguity resolution (Vitello et al., 2014), rather than to the specific linguistic tasks at hand (i.e., whether semantic, phonological, or syntactic; see Snyder et al., 2007; January et al., 2009; Rodd et al., 2010; Hsu et al., 2017; Novick et al., 2009, for evidence in brain-damaged patients; but see Santi & Grodzinsky, 2007 for conflicting results).

A subset of bilateral frontal and parietal cortices have been identified as involved in different types of cognitively challenging tasks (cf. Duncan & Owen, 2000; Duncan, 2010, 2013). This set of brain areas has collectively been labelled the ‘Multiple-Demand System’ (MDS), and includes the cortex surrounding the posterior inferior frontal sulcus (LIFC), anterior insular cortex, premotor cortex, dorsolateral prefrontal cortex, anterior cingulate cortex, pre-supplementary motor area, and the cortex surrounding the intraparietal sulcus. A defining functional characteristic of this network is its consistent activation/engagement during cognitive or executive control tasks. More specifically, these regions are sensitive to cognitive demands, namely the level of difficulty across many different domains, such as perception, language, memory, response selection, response inhibition, problem solving, task novelty and so on, typically showing increased activity in more challenging conditions (Fedorenko et al., 2013; Woolgar et al., 2013).

Building on this approach, in a recent paper Fedorenko and colleagues have investigated whether activity in the LIFC is language-specific or domain-general in terms of the functional properties exhibited by the MDS (Fedorenko et al., 2012). Using a linguistic (sentence reading) vs. a non-linguistic task (non-words reading) they identified sub-regions within the LIFC which were either sensitive or insensitive to linguistic processing. Subsequently, two sub-regions were investigated during the performance of six different cognitive tasks (arithmetic addition, spatial/verbal working memory, Stroop task, and two versions of the multisource interference task), each of which included an ‘easier’ and a ‘harder’ condition. Their results showed that Broca’s area contained two functionally distinct sub-regions lying side by side. A first sub-region (located in the triangular part of the LIFC), was highly responsive to the processing of linguistic material, but showed little or no response to cognitive tasks and/or the degree of cognitive challenge. A second sub-region (surrounding the first one), showed instead little or no response to linguistic processing, but was extremely sensitive to cognitive tasks (irrespective of the stimulus material used), and more active in harder rather than easier conditions. The authors concluded that Broca’s area is not a homogenous functional unit. Instead, within Broca’s area there are both language-specific and domain-general units.

Related studies from the same group have reported consistent results, showing that other nodes within the language network (e.g., superior temporal and inferior parietal cortices) do not show any sensitivity to cognitive demand/difficulty (Fedorenko et al., 2011), whereas nodes within the MDS do exhibit such a sensitivity (Fedorenko et al., 2013). Furthermore, the two networks (language areas vs. MDS) show a dissociation in functional connectivity (i.e., internal coherence) and a reciprocal lack of correlation (Blank et al., 2014). These studies have provided us with very valuable contributions to understand how the LIFC and MDS work. However, like all studies, they also have a number of limitations. First, the cognitive tasks adopted (as well as the harder and easier conditions) were not designed to be directly comparable to one another (as acknowledged by Fedorenko et al., 2013). Second, cognitive challenge in the linguistic task was not manipulated, so it is unclear how activity in the triangular part of the LIFC is modulated by linguistic challenge. Third, they made use of a non-standard, subject-based analytical approach (Fedorenko et al., 2010). Although this approach has the benefit of taking into account individual differences in functional anatomy, it makes it difficult to draw inferences at the population level.

To address these limitations, we designed a double-blind randomised crossover functional neuroimaging (fMRI) study to investigate which parts of the LIFC are engaged in a domain-specific manner, and which ones are engaged in a domain-general manner, i.e. to identify sub-regions within the LIFC whose activity is modulated according to a clear functional rule (i.e., domain-specificity vs. -generality). We developed two tasks: one linguistic (picture Naming) and one non-linguistic (size Judgment) with two difficulty levels (High vs. Low) carefully matched in terms of: stimulus material, experimental conditions, output demand, and behavioural performance (see below for details).

This enabled us to first delineate the neural correlates associated with the specific cognitive processes central to each task, and ask whether they recruit the LIFC to a similar or differential degree (i.e., testing domain-specificity vs. domain-generality). According to the aforementioned theoretical standpoints, for domain-specific sub-regions within the LIFC (i.e., hubs of the ‘Language Network’), which are functionally specialised (i.e., ‘modular’) for linguistic processing (e.g., Fedorenko et al., 2012), we should predict: i) little-to-no overlapping activation between the two tasks; and/or ii) different activation patterns for the two tasks in sub-regions modulated by different functional rules (e.g., greater BOLD response for naming vs. judgment, and vice-versa). Conversely, domain-general sub-regions should support both tasks performance with increased recruitment reflecting increasing cognitive challenge irrespective of the nature of the stimuli at hand (e.g., Duncan, 2010). In such sub-regions we should predict (at least partial) overlapping activation between the two tasks, and/or similar activation patterns (i.e., BOLD response profiles) across both tasks.

Second, we could cleanly isolate the neural activation patterns associated with cognitive demand (namely difficulty), across both tasks while controlling for stimulus type. This allowed us to investigate whether, and where, cognitive demand (High-challenge) increases activity in the LIFC to a greater extent than Low-challenge (i.e., testing domain-generality). For domain-specific sub-regions, we should predict either: i) different patterns of sensitivity to cognitive challenge modulations for the two tasks (e.g., increased activity for High->Low-challenge for Naming and Judgment in different sub-regions); or ii) no increased BOLD response in the Judgment task. In domain-general sub-regions, we should predict comparable response patterns to cognitive challenge modulations across both tasks (e.g., increased activity for High->Low-challenge in both tasks in the same sub-regions).

Additionally, we delivered two types of transcranial direct current stimulation (tDCS) to the LIFC concurrently with the fMRI study (Anodal vs. Sham). In this way, we were able to investigate whether and how neuromodulation of the LIFC affects on-line brain and behavioural performance for specific cognitive processes (linguistic vs. non-linguistic) for each task, and general cognitive demand, namely difficulty (Low vs. High) across both tasks (i.e., testing the contribution of the LIFC to specificity/generality). Anodal tDCS delivered to the LIFC has been shown to reduce both reaction times (RTs) and BOLD response within the LIFC during a spoken naming fMRI task (Holland et al., 2011). This was interpreted as brain and behavioural priming by tDCS. The electrode covers a relatively large area and is supposed to stimulate both domain-specific and domain-general sub-regions. However, from a domain-specific perspective, we should predict neuromodulation to result in different behavioural and brain effects across tasks. For example, facilitation of the linguistic (but not non-linguistic) task, or behavioural effects in both tasks but different underlying neural activation patterns in different sub-regions of the LIFC for each task (i.e., different neural interactions). Conversely, from a domain-general perspective, we should predict tDCS to result in significant behavioural and brain effects across both tasks and within the same LIFC sub-region(s). For example, anodal tDCS facilitating behavioural responses (reduced RTs) across both tasks with a corresponding modulation of neural activation within the same hub.

Viewed broadly, our study aimed – for the first time – to compare the neurocognitive architecture of spoken language processing across domains and tasks by matching task and stimulus characteristics. Functional neuroimaging enabled us to characterise, within subjects, the common and dissociable neural correlates underlying multiple levels of two demanding tasks (one requiring spoken language). While neuromodulation of the LIFC would allow us to directly test whether its contribution to spoken language processing is domain-specific (i.e., language preferential) relying on independent components (LIFC sub-regions), domain-general (i.e., shared resources across both our cognitively challenging tasks), or perhaps a more nuanced picture of both.

## METHODS

### Participants

This study is part of a larger research project about anomia rehabilitation in people who suffered left hemisphere stroke. In this framework, we recruited a cohort of healthy controls who will be later compared with aphasic stroke patients. Here, we report the data of 17 healthy right-handed native English speakers (6 M, mean age: 69±9), who took part in the study. All had normal hearing, normal or corrected-to-normal visual acuity, no history of neurological or psychiatric disease, and no contraindications to MR scanning. All participants gave written informed consent to participate in the study, which was approved by the Central London Research Ethics Committee and conducted in accordance with the ethical principles stated by the Declaration of Helsinki.

### Stimuli and experimental conditions

Each experimental trial consisted of the simultaneous presentation of an auditory cue associated with the picture of a concrete object (cf. Figure 1A). A list of 480 target words was drawn from the IPNP database (n=220), and from the MRC Psycholinguistic Database (n=260; http://websites.psychology.uwa.edu.au/school/MRCDatabase/mrc2.html; Coltheart, 1981). All object names were monosyllabic words and consonant-vowel-consonant (CVC) in terms of phonological structure. Auditory cues consisted of either the initial phoneme of a target word, or a noise control. To generate the auditory cues, each target word was digitally recorded (at 44.1 kHz) from a male native English speaker in a soundproof room, then cropped at the offset of the vowel to form the initial phoneme cue (e.g., /bp/ for ‘box’). Noise control cues were generated by noise vocoding the initial phoneme cues. This was performed utilizing the technique described by Shannon et al. (1995), using custom Matlab scripts (cf. Evans & Davis, 2015). Accordingly, the frequency range of 30-6000 Hz was divided into a single channel. The amplitude envelope was extracted by half-wave rectifying the signal and applying a low-pass filter with a cut-off of 30 Hz, to remove pitch synchronous oscillations. This envelope was used to amplitude modulate band-pass filtered white noise in the same frequency range as the source. This generated an acoustic signal with a temporal and spectral profile similar to the original speech, but not intelligible. Initial phoneme and noise control cues were matched for auditory duration. Visual stimuli consisted of 480 black and white line drawings of concrete objects, partly derived from the International Picture Naming Project (IPNP; Szekely et al., 2004; http://crl.ucsd.edu/experiments/ipnp/index.html), and the remainder found on the internet (with similar style/figurative features as the IPNP items).

**Figure 1 –.**
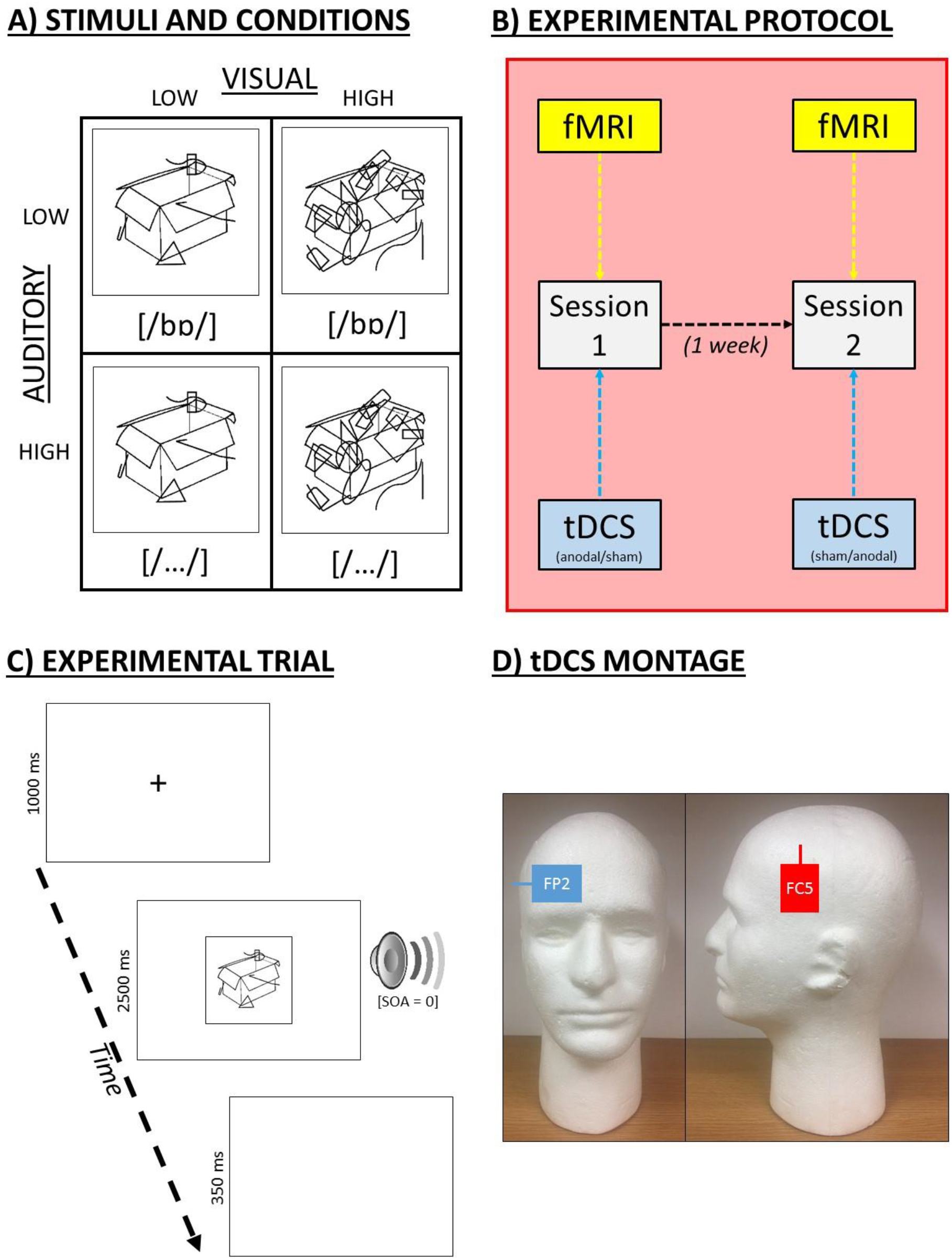
Experiment. A) Examples of stimuli and experimental conditions. Stimuli always consisted of a picture presented concurrently with an auditory cue, and cognitive challenge was varied orthogonally in two sensory modalities (i.e., auditory and visual) at a time. Here, an example item (box) is shown in auditory Low- and High-challenge conditions (initial cue vs. noise, respectively), accompanied by visual Low- and High-challenge conditions (5 vs. 15 masking elements overlapped, respectively). B) Experimental protocol showing the two concurrent fMRI-tDCS sessions. C) Example of an experimental trial. Concurrent delivery of the auditory and visual stimuli were preceded by an alerting fixation cross, and followed by a blank screen. D) tDCS montage. Example positioning of the anodal (red; FC5) and cathodal (blue; FP2) electrodes onto the head. Legend: SOA = stimulus-onset asynchrony.

Our experimental conditions were designed in order to manipulate cognitive challenge orthogonally in the auditory and visual modalities at the same time. Aurally, each picture was presented simultaneously with an auditory cue in two experimental conditions: i) Low-challenge (initial phoneme); or ii) High-challenge (noise-vocoded control). Visually, pictures were presented with a variable amount of visual noise overlapped (i.e., masking elements made up of black squiggly lines and/or geometrical shapes), in two experimental conditions: i) Low-challenge (5 masking elements); or ii) High-challenge (15 masking elements; cf. Figure 1A). Both manipulations had the effect of making an object more ambiguous, and therefore increasing cognitive challenge to identify an object identity. Items were assigned to the various experimental conditions in such a way that average psycholinguistic features (e.g., frequency, concreteness, imageability, initial phoneme, etc.) were balanced across conditions, and assignment was counterbalanced across subjects and sessions (see below).

### Tasks and procedure

Subjects performed two fMRI-tDCS sessions (either Anodal or Sham tDCS on each occasion, see below) one week apart (cf. Figure 1B), with the order counterbalanced across subjects. In each session, subjects were required to perform two tasks in different functional runs: i) a picture Naming task; and ii) a size Judgment task. Subjects performed two runs of Naming and two runs of Judgment per session, and the sequence of tasks in the four functional runs was counterbalanced both between subjects and sessions. In the Naming task, subjects had to name each target picture as quickly and as accurately as possible. In the Judgment task, they had to determine (yes/no spoken responses) whether the size of each object depicted would fit inside a microwave oven. This type of decision was required because – contrary to decisions such as living vs. non-living, natural vs. man-made, or indoor vs. outdoor – the answer is not already available in semantic memory, i.e. it requires new item-specific processing in real time. In terms of cognitive processes involved, both tasks required object identification, decision making, and a vocal response. However, while the Naming task necessarily relies upon lexical retrieval, this is not the case with the Judgment task.

Each visual stimulus was displayed for 2500 ms, preceded by a 1000 ms alerting fixation cross and followed by a blank screen for 350 ms (see Figure 1C). Auditory cues were presented simultaneously with each picture (Stimulus-Onset-Asynchrony=0 ms). Trials were presented in mini-blocks of six stimuli (belonging to different conditions), separated by fixation-only rest periods of 7700 ms in order to optimize the timing of the experiment for the BOLD response (Henson, 2006). To vary the spatiotemporal synchrony between the trial structure and the image acquisition the inter-trial interval was set to 3850 ms to jitter the onset of each trial across acquired brain volumes.

Overt spoken responses were recorded online using a dual-channel, noise-cancelling fibre optical microphone system (FOMRI III; http://www.optoacoustics.com), and reviewed offline to determine trial-specific reaction times (RTs) for each subject. Auditory cues were delivered via MR-compatible headphones (MR Confon, Magdeburg, Germany; www.mr-confon.de). The order of experimental conditions was pseudo-randomized within a functional run (i.e., avoiding more than three trials of the same condition in a row). On each session, all subjects underwent a short training period, before entering the MR scanner, to become familiar with the tasks and to practice how to speak in a soft voice to minimise motion in the scanner. Stimuli used during the training were not used during the fMRI session.

### Transcranial direct current stimulation (tDCS)

tDCS was delivered during the fMRI experiment by using an MR-compatible stimulation system (neuroConn; https://www.neurocaregroup.com/dc_stimulator_mr.html) via a pair of MR-compatible leads and rectangular rubber electrodes (5×7 cm), allowing for a current density of 0.057 mA/cm^2^ (cf. Holland et al., 2011). For all participants, the anode was placed over the LIFC (equivalent to position FC5 in a 10-20 EEG nomenclature; cf. Figure 1D), and the cathode placed over the contralateral frontopolar cortex (FP2). Both electrodes and the sites on the scalp where the electrodes were placed were covered with EEG conductive paste to ensure a flush and comfortable fit between the electrode surface and the scalp. Electrodes were secured to the head using 3M Coban elastic wrap bandage and placed in adherence with the manufacturer’s MR safety guidelines. Care was taken in connecting the leads backward along the centre of the scanner bore to minimize the possibility of radio frequency-induced heating, and to ensure that any gradient switching-induced AC currents were well below the level that might cause stimulation. The stimulator was placed outside the Faraday cage of the scanner, and the stimulating current was fed to the participant through two stages of radio frequency filtration to prevent interference being picked up by the scanner.

A scanner pulse triggered the onset of the stimulation at a given slice in the acquisition sequence. The current was increased slowly during the first 15 sec to the desired stimulation threshold (2 mA), termed the ramp-up phase. A constant direct current (2 mA) was delivered for 20 min. At the end of the stimulation period, the current was decreased to 0 mA over 1 sec (ramp-down). For sham stimulation, the ramp-up phase was followed by 15 sec of 2 mA stimulation, which was immediately followed by a 1 sec ramp-down phase. This active sham protocol resulted in a more efficient blinding process.

tDCS stimulation was conducted in a double-blind paradigm. Both stimulation and sham protocols produced sensations of comparable quality (a mild tingling, typically under the electrode placed over the contralateral orbital/frontopolar edge). Participants habituated to it quickly and reported minimal discomfort with no adverse sensations, phosphenes, or analogous effects during anodal and sham tDCS stimulation runs. Four out of 17 subjects reported detecting a difference between the two sessions. However, they could not identify reliably which was the sham or the anodal stimulation session, i.e. their responses were at chance level. The position of the anode and cathode electrodes for each subject was recorded and reproduced across scanning sessions.

### Imaging acquisition and analysis

Whole-brain imaging was performed on a 3T Siemens TIM-Trio system (Siemens, Erlangen, Germany) at the Wellcome Centre for Human Neuroimaging. T2*-weighted echo-planar images (EPI) with BOLD contrast were acquired using a 12-channel head coil. Imaging was optimised for BOLD sensitivity in the inferior frontal cortex (Weiskopf et al., 2006). Each EPI volume comprised 48 axial slices with sequential ascending acquisition, slice thickness=2.5 mm, inter-slice gap=0.5 mm, in-plane resolution=3×3 mm^2^. Volumes were acquired with a TR=3360 ms, and the first six volumes of each session were discarded to ensure a steady state had been reached. In each session, a total of 195 volume images (189 volumes of interest and 6 dummy scans) were acquired in each of four consecutive runs, each lasting approximately 11 min. Prior to the first functional run of each scanning session, a dual gradient-echo based field map was acquired for each subject for later B0 field distortion correction of functional images. The same scanner and hardware were used for the acquisition of all images. Functional data were pre-processed and analysed using Statistical Parametric Mapping software (SPM12; www.fil.ion.ucl.ac.uk/spm) running under Matlab 2015a (MathWorks, Natick, MA). All volumes of interest from each subject were realigned and unwarped, using session- and subject-specific voxel displacement maps (Hutton et al., 2002). The functional images were then co-registered with the structural image, spatial normalisation parameters were estimated using this latter and applied to functional images. Finally, functional data were spatially smoothed with an 8 mm FWHM isotropic Gaussian kernel to account for residual misalignment after spatial normalization and the application of Gaussian Random Field Theory for corrected statistical inference. To remove any low-frequency drifts, data were high-pass filtered using a set of discrete cosine functions with a cut-off period of 128 sec.

Statistical analyses were first performed in a subject-specific fashion. Eight conditions per session (i.e., 2 Tasks x 2 Visual Challenge levels x 2 Auditory Challenge levels) were modelled separately as events convolved with the SPM canonical haemodynamic response function (HRF). We used the presentation of the concurrent auditory cue/picture as the onset of the event. Movement realignment parameters were included as covariates of no interest. The resulting stimulus-specific parameter estimates were calculated for all brain voxels using the General Linear Model. At the second level, 16 conditions of interest were modelled (2 Tasks x 2 Visual Challenge levels x 2 Auditory Challenge levels x 2 tDCS stimulation conditions), modelling subjects as a random factor. Significance threshold for all reported results was set to p<0.05 FWE-corrected for multiple comparisons either across the whole-brain, or within *a priori* hypothesised regions-of-interest (ROIs) within the LIFC (i.e., when a small-volume-correction was applied, see below for details). Anatomical labelling was determined by using the Automated Anatomical Labelling atlas (AAL; Tzourio-Mazoyer et al., 2002).

## RESULTS

### Behavioural results

We performed a 2 x 2 x 2 x 2 repeated-measure ANOVA on reaction times (RTs) of all responses (cf. Supplementary Material) with ***Task*** (Naming, Judgment), ***Auditory Challenge*** (Low, High), ***Visual Challenge*** (Low, High), and ***tDCS*** (Anodal, Sham) as within-subject variables (Figure 2A). Significance threshold for reported results was set to p<0.05 throughout (see Table 1 for ANOVA results).

**Figure 2 –.**
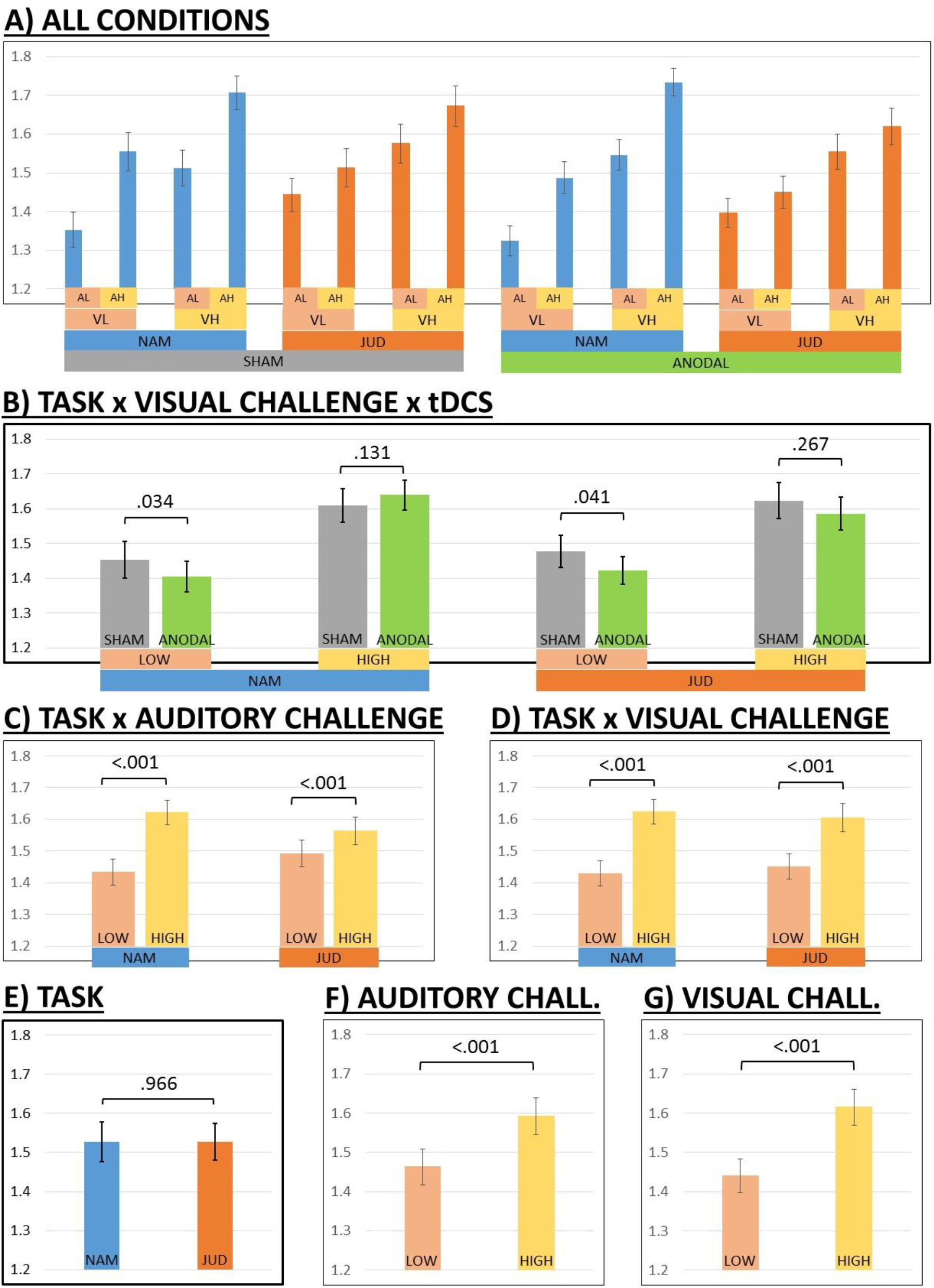
Behavioural results (reaction times). A) Bars showing all experimental conditions. B) Task x Visual challenge x tDCS interaction (averaged across Auditory challenge). C) Task x Auditory challenge interaction (averaged across Visual challenge). D) Task x Visual challenge interaction (averaged across Auditory challenge). E) Main effect of Task (averaged across Auditory challenge, Visual challenge, and tDCS). F) Main effect of Auditory challenge (averaged across tasks, Visual challenge, and tDCS). G) Main effect of Visual challenge (averaged across tasks, Auditory challenge, and tDCS). All post-hoc comparisons are computed two-tailed. Legend: NAM = Naming; JUD = Judgment; AL = Auditory Low-challenge; HL = Auditory High-challenge; VL = Visual Low-challenge; VH = Visual High-challenge; LOW = Low-challenge; HIGH = High-challenge. Y axis in plots shows reaction time (RTs) in seconds.

**Table 1 -.**
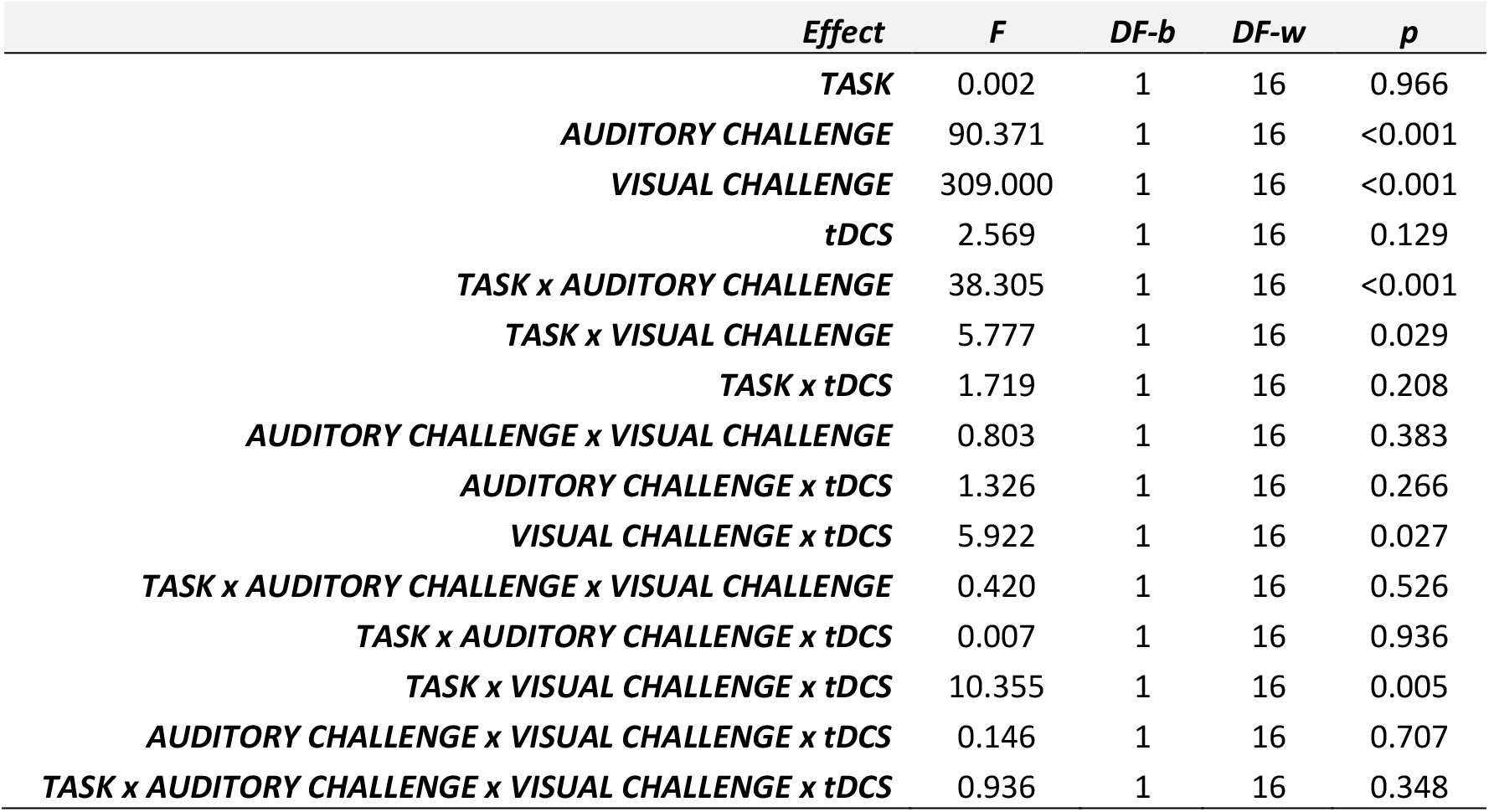
Behavioural results (ANOVA on reaction times). Legend: F = F-test; DF-b = degrees of freedom between; DF-w = degrees of freedom within; p = p-values

| *Effect* | *F* | *DF-b* | *DF-w* | *p* |
| --- | --- | --- | --- | --- |
| **TASK** | 0.002 | 1 | 16 | 0.966 |
| **AUDITORY CHALLENGE** | 90.371 | 1 | 16 | <0.001 |
| **VISUAL CHALLENGE** | 309.000 | 1 | 16 | <0.001 |
| **tDCS** | 2.569 | 1 | 16 | 0.129 |
| **TASK x AUDITORY CHALLENGE** | 38.305 | 1 | 16 | <0.001 |
| **TASK x VISUAL CHALLENGE** | 5.777 | 1 | 16 | 0.029 |
| **TASK x tDCS** | 1.719 | 1 | 16 | 0.208 |
| **AUDITORY CHALLENGE x VISUAL CHALLENGE** | 0.803 | 1 | 16 | 0.383 |
| **AUDITORY CHALLENGE x tDCS** | 1.326 | 1 | 16 | 0.266 |
| **VISUAL CHALLENGE x tDCS** | 5.922 | 1 | 16 | 0.027 |
| **TASK x AUDITORY CHALLENGE x VISUAL CHALLENGE** | 0.420 | 1 | 16 | 0.526 |
| **TASK x AUDITORY CHALLENGE x tDCS** | 0.007 | 1 | 16 | 0.936 |
| **TASK x VISUAL CHALLENGE x tDCS** | 10.355 | 1 | 16 | 0.005 |
| **AUDITORY CHALLENGE x VISUAL CHALLENGE x tDCS** | 0.146 | 1 | 16 | 0.707 |
| **TASK x AUDITORY CHALLENGE x VISUAL CHALLENGE x tDCS** | 0.936 | 1 | 16 | 0.348 |

### Task

Critically, RT data showed no significant main effect of ***Task***, that is, the two tasks were behaviourally matched overall (Figure 2E).

### Cognitive challenge

#### INTERACTIONS

We found both a significant ***Task* x *Auditory Challenge*** and a ***Task* x *Visual Challenge*** interaction (Figure 2C-D). This implied that the difference between High-challenge and Low-challenge was significantly larger in the Naming than in the Judgment task.

#### MAIN EFFECTS

There was a significant main effect of both ***Auditory Challenge*** and ***Visual Challenge*.** As predicted, High-challenge conditions resulted in significantly slower RTs with respect to Low-challenge (Figure 2F-G) conditions.

### tDCS

#### INTERACTIONS

There was a significant three-way ***Task x Visual Challenge x tDCS*** interaction. This showed that – with respect to Sham tDCS – Anodal tDCS significantly reduced RTs in Low-challenge conditions across both tasks, whereas in High-challenge conditions there was an opposite non-significant effect of tDCS: a trend to increased RTs (slower responses) in the Naming task and a trend to reduced RTs (faster responses) in the Judgment task (cf. Figure 2B).

This result was consistent with a significant ***Visual Challenge x tDCS*** interaction, where – with respect to Sham tDCS – Anodal tDCS significantly reduced RTs for Low-challenge (p=0.013) but not for High-challenge (p=0.878) conditions, irrespective of task.

#### MAIN EFFECTS

Finally, we found no significant main effect of ***tDCS***. However, to test whether previous results from our group could be replicated, we ran a subsidiary ANOVA only on the Low-challenge visual condition, that is, the most similar condition to the one used in our previous picture naming study (i.e., no visual masking; cf. Holland et al., 2011). This revealed a significant main effect of ***tDCS*** (p=0.014), showing that Anodal tDCS significantly reduced RTs both in the Naming and in the Judgment task with respect to Sham tDCS (cf. three-way interaction above and Low-challenge bars in Figure 2B).

In summary, behavioural results showed that: i) the two tasks were equally demanding; ii) the challenge modulation was effective in both modalities, in the predicted direction (High>Low), and with a wider range in the Naming task; iii) Anodal tDCS significantly facilitated visually Low-challenge (but not High-challenge) conditions in both tasks.

### Imaging results

#### Task

The Naming and the Judgment tasks showed the recruitment of a widespread, largely overlapping neural network. This was revealed by the main effect of experiment (***Naming&Judgment>Rest***; Figure 3A) and confirmed by the contrasts ***Naming>Rest*** and ***Judgment>Rest*** (cf. Supplemental Figure S1A-B). It comprised bilateral inferior frontal cortices (although most prominently on the left, including the opercular, triangular, and orbital parts), most nodes of the MDS such as anterior insular cortices, premotor cortices, pre-supplementary motor areas, dorsal anterior cingulate cortices, posterior parietal cortices, as well as visual (occipital, occipito-temporal, and occipito-parietal) and auditory (superior temporal, middle temporal, temporal polar, and inferior parietal) cortices (Table 2).

**Figure 3 –.**
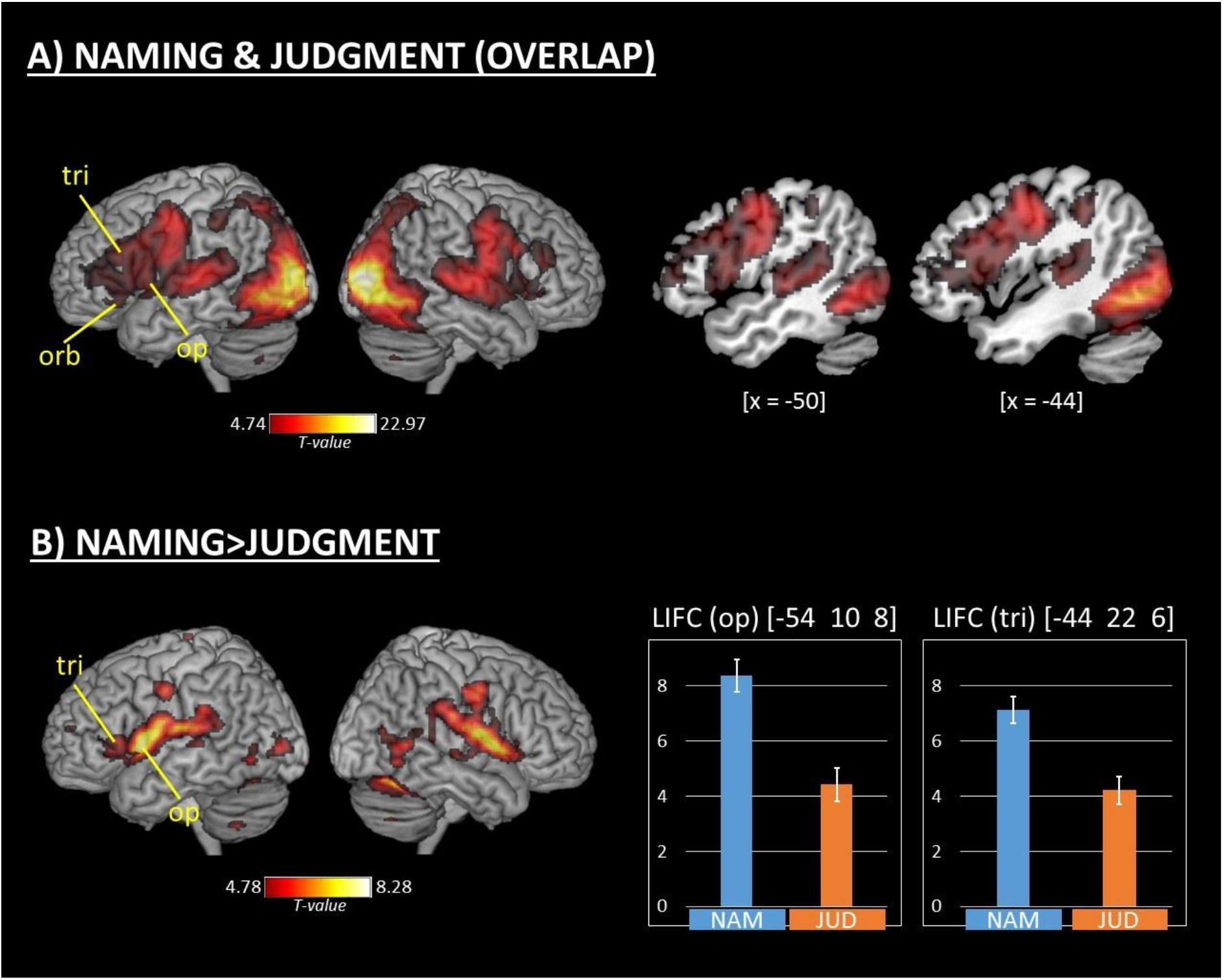
Tasks-related activations. A) Areas activated by both Naming and Judgment tasks (i.e., main effect of experiment, or Naming & Judgment > Rest; cf. also Supplementary Figure S1A-B). B) Areas activated more in the Naming than in the Judgment task (i.e., main effect of Task). Legend: op = opercular part of the LIFC; tri = triangular part of the LIFC; orb = orbital part of the LIFC; NAM = Naming; JUD = Judgment. Y axis in plots shows effect size of BOLD response in arbitrary units.

**Table 2 -.**
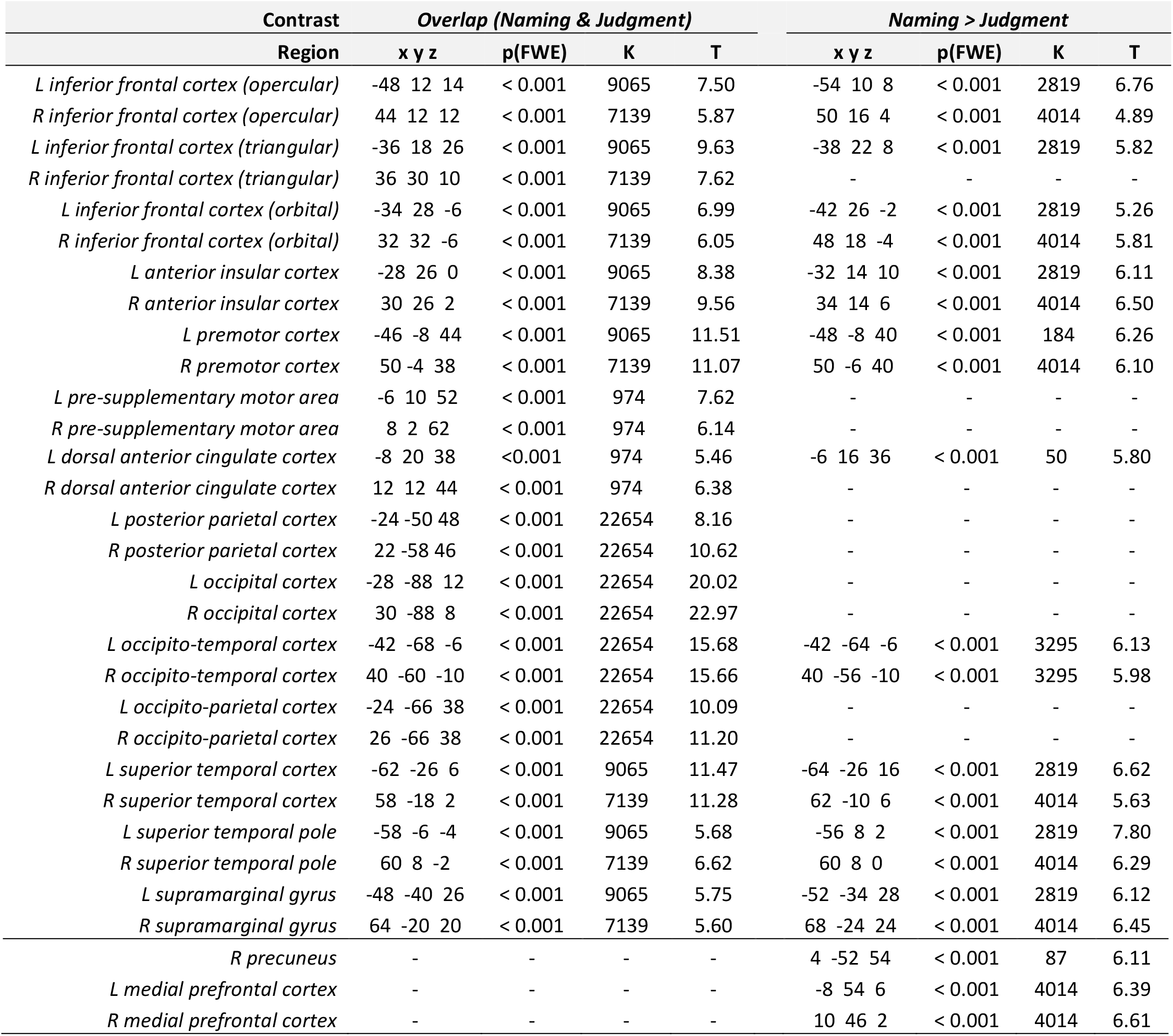
Tasks-related activations. Legend: R = right; L = left; x y z = MNI coordinates; K = cluster size; T = t-scores; p(FWE) = family-wise error corrected p-values

A direct comparison between the two tasks (i.e., main effect of Task: ***Naming>Judgment***) showed a subset of areas within this common network significantly more activated in the Naming than in the Judgment task (Figure 3B). This included bilateral portions of the inferior frontal cortices (more widespread on the left, including bilateral opercular and orbital, and left triangular parts), anterior insular cortices, premotor cortices, superior temporal and inferior parietal cortices, temporal poles, as well as visual (occipital and occipito-temporal) cortices, plus the left dorsal anterior cingulate cortex. Outside the common network, significant activations were found in bilateral ventral medial prefrontal cortices, and in the right precuneus (Table 2). The reverse contrast (***Judgment>Naming***) showed no significant results.

### Cognitive challenge

#### INTERACTIONS

Mirroring our behavioural results, there were significant ***Task x Auditory Challenge*** and ***Task x Visual Challenge*** interactions. In the auditory modality, effects were found in three LIFC clusters (opercular, triangular, and orbital parts) as well as in the pre-supplementary motor area (Figure 4A, Table 3). Critically, activation patterns in all these nodes showed that – as compared with Low-challenge conditions – High-challenge conditions were associated with increased BOLD response in the Naming task, but with decreased BOLD response in the Judgment task (cf. plots in Figure 4A), i.e. a differential activation pattern across conditions in the two tasks in the same sub-regions. In the visual modality, a significant cluster was identified in the right inferior occipito-temporal cortex (Figure 3B, Table 3), also showing that – as compared with Low-challenge conditions – High-challenge conditions were associated with increased BOLD response in the Naming task, but with decreased BOLD response in the Judgment task.

**Figure 4 –.**
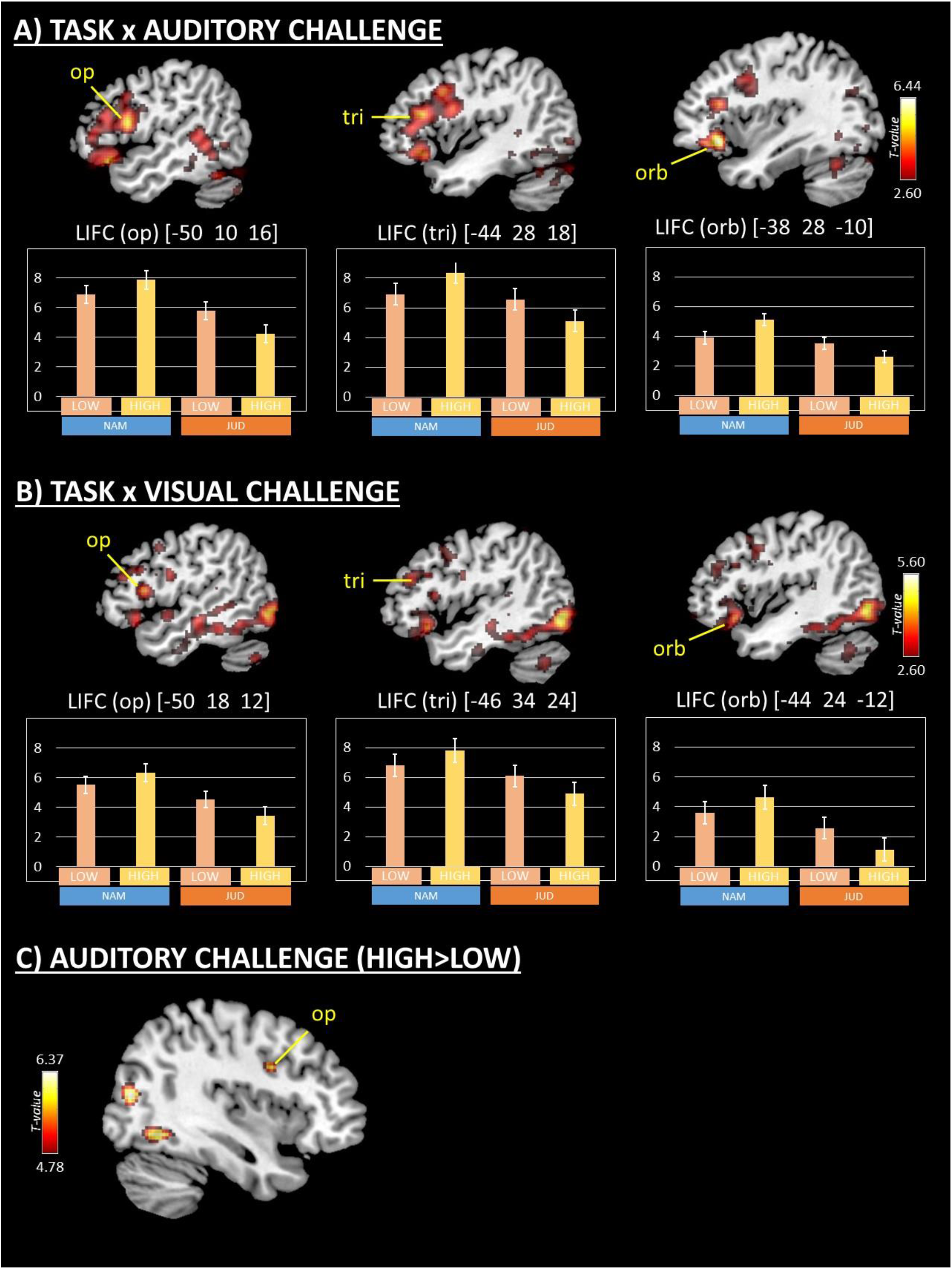
Cognitive challenge-related activations in the LIFC. A) Areas activated by the Task x Auditory challenge interaction. B) Areas activated by the Task x Visual challenge interaction. C) Areas associated with Auditory challenge (High->Low-challenge). In (A) and (B) activations are shown at p <0.005-unc. for display purposes. Peaks shown in (A) are significant at whole-brain level (p<0.05 FWE-corr.). Peaks shown in (B) survive small-volume correction (cf. main text). In (C) activations are shown at p<0.05 FWE-corr. Legend: op = opercular part of the LIFC; tri = triangular part of the LIFC; orb = orbital part of the LIFC; NAM = Naming; JUD = Judgment; LOW = Low-challenge; HIGH = High-challenge. Y axis in plots shows effect size of BOLD response in arbitrary units.

**Table 3 -.**
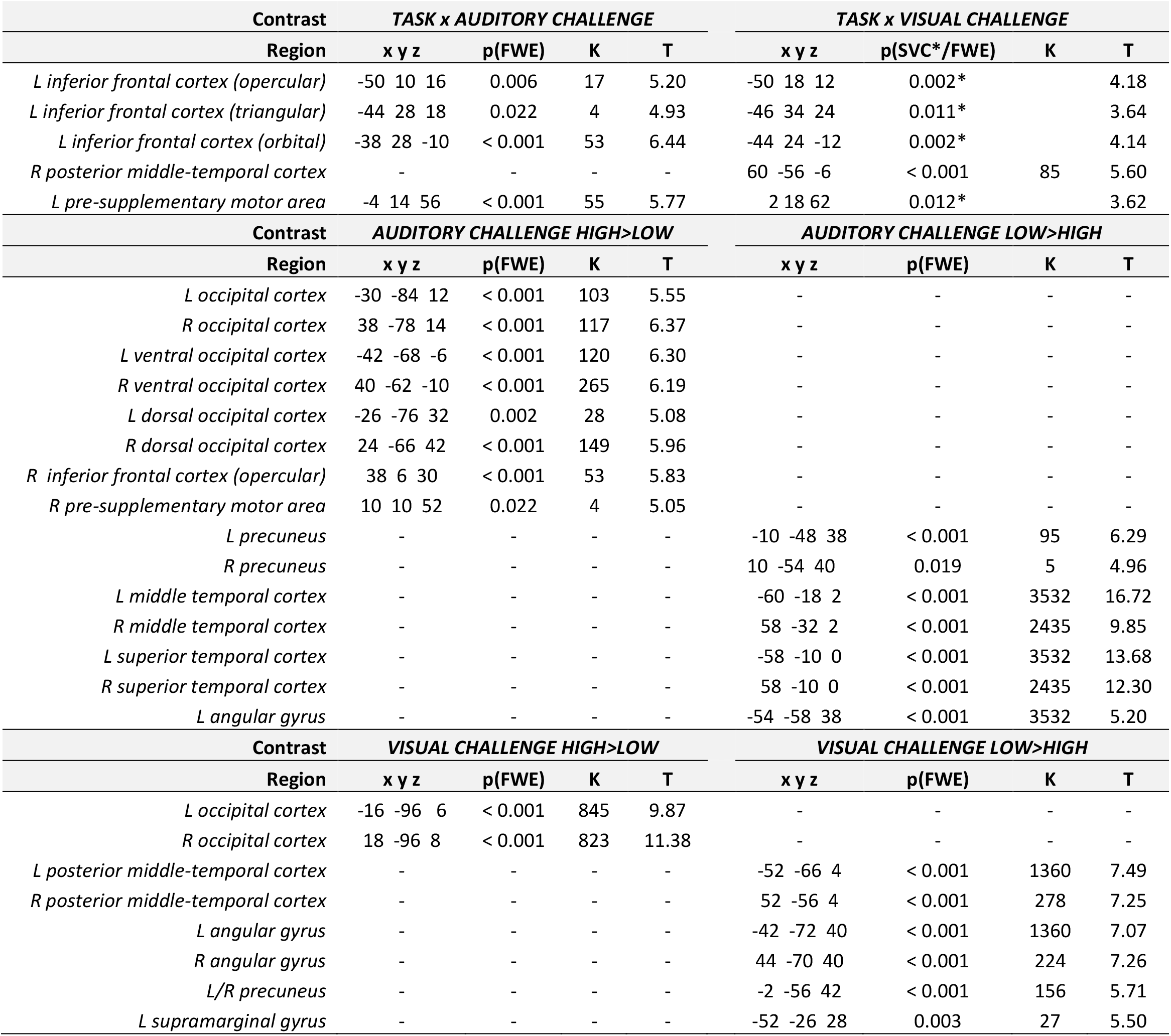
Cognitive challenge-related activations. Legend: R = right; L = left; x y z = MNI coordinates; K = cluster size; T = t-scores; p(FWE) = family-wise error corrected p-values; p(SVC*) = small-volume-corrected p-values

In the visual modality, activations in the LIFC did not survive whole-brain correction. However, in order to test whether there was any sub-threshold cluster in the ***Task x Visual Challenge*** contrast showing an activation pattern such as the one identified by the ***Task x Auditory challenge***, we performed a region-of-interest (ROI) analysis. Accordingly, a small-volume-correction within three spheres of 10 mm radius centred in the peaks of the four clusters identified by the ***Task x Auditory Challenge*** contrast (i.e., orthogonal to the one tested; cf. coordinates in the top-left panel of Table 3) was applied. This analysis revealed significant visual modulatory effects in all sub-regions (see top-right panel of Table 3), showing exactly the same activation pattern as in the auditory modality (i.e., ***Task x Auditory challenge*** interaction), in exactly the same neural network (three clusters in the LIFC; cf. plots in Figure 4B).

#### MAIN EFFECTS

The main effect of ***Auditory challenge*** was a significant modulation of bilateral visual and auditory activations (Supplemental Figure S1C, Table 3). In the auditory modality, the contrast ***High-challenge>Low-challenge*** revealed increased activity in associative (occipital and occipito-temporal) visual regions, at the border between the right premotor cortex and the right inferior frontal cortex (opercular part; see Figure 4C) and in the pre-supplementary motor area. The contrast ***Low-challenge>High-challenge*** showed increased activity in bilateral auditory (superior temporal, extending into the middle temporal) cortices, and the precuneus.

The main effect of ***Visual challenge*** was a significant modulation of activity in bilateral visual cortices (Supplemental Figure S1D, Table 3). In the visual modality, the contrast ***High-challenge>Low-challenge*** showed increased activity in primary and secondary (occipital) visual areas, whereas the contrast ***Low-challenge>High-challenge*** showed increased activity in associative (occipito-temporal and occipito-parietal) visual areas (extending into the inferior parietal cortex), and the precuneus.

Critically, neither the main effect of ***Auditory challenge*** nor the main effect of ***Visual challenge*** showed significant modulations of brain activity in the LIFC.

#### MAIN EFFECTS AT A LOWER THRESHOLD

The MDS theory predicts that increasing cognitive challenge is associated with increased activity within the MDS, and indeed we found that both the Naming and the Judgment tasks significantly recruited several classical MDS nodes (cf. overlap in Figure 3A and Table 2). We then used this same overlap map as a masking ROI for the contrasts ***High-challenge>Low-challenge*** to qualitatively check whether challenge-related modulations in the MDS occurred sub-threshold (p<0.005-unc.). This revealed activity in several MDS nodes: bilateral anterior insular cortices, premotor cortices, dorsal anterior cingulate cortices, pre-supplementary motor areas, and posterior parietal cortices in both modalities (cf. Figure 5 and Supplementary Table ST1). Critically, none of these regions exhibited an activation pattern similar to the one identified in the LIFC. Indeed – in line with the MDS theory predictions – all these regions show an increased activity in High-challenge conditions vs. Low-challenge conditions in both tasks (although often with a larger observed effect in the Naming task).

**Figure 5 –.**
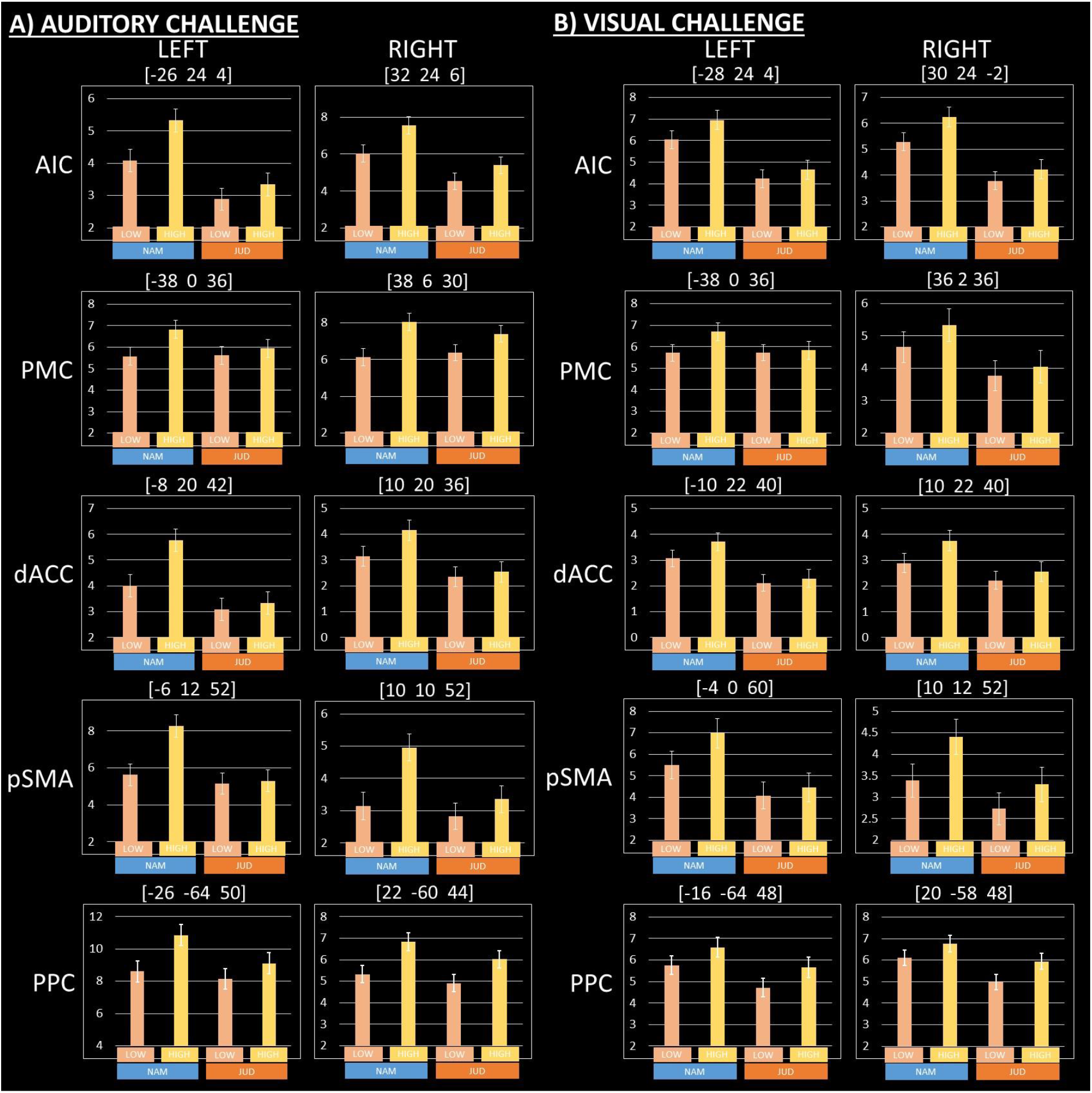
Cognitive challenge-related activations in other Multiple-Demand System (MDS) nodes, showing the same activation patterns in the two tasks. A) Modulation by Auditory challenge. B) Modulation by Visual challenge. Number in square brackets report MNI coordinates (x y z) of the peak plotted (cf. Supplementary Table ST2). Legend: NAM = Naming; JUD = Judgment; LOW = Low-challenge; HIGH = High-challenge; AIC = anterior insular cortex; PMC = premotor cortex; dACC = dorsal anterior cingulate cortex; pSMA = pre-supplementary motor area; PPC = posterior parietal cortex. Y axis in plots shows effect size of BOLD response in arbitrary units.

### tDCS

#### INTERACTIONS

We did not find any significant interaction with tDCS at the chosen statistical threshold (p<0.05 FWE-corrected). However, mirroring our behavioural results, we found a sub-threshold ***Task x Visual Challenge x tDCS*** interaction in the LIFC (triangular part; see Figure 6). This region was located in the same area identified by a previous study of ours showing that Anodal tDCS – delivered to the LIFC, consistent electrode montage as in the present study – was associated with reduced activity in this area during speech production (Holland et al., 2011). Hence, in order to formally test this sub-threshold activation, we performed an ROI analysis. Accordingly, a small-volume-correction within a sphere of 10 mm radius centred in the triangular part of the LIFC (x y z = −48 32 19; cf. Holland et al., 2011) was applied. This analysis showed a significant modulatory effect (x y z = −52 36 10; T = 3.28; p = 0.032), replicating our previous naming results. Here, the activation pattern showed that – with respect to Sham tDCS – Anodal tDCS was associated with a greater reduction in BOLD response in Low-challenge than High-challenge conditions in the Naming task, and vice-versa in the Judgment task (i.e., greater reduction in BOLD response in High-challenge than Low-challenge conditions; cf. plot in Figure 6).

**Figure 6 –.**
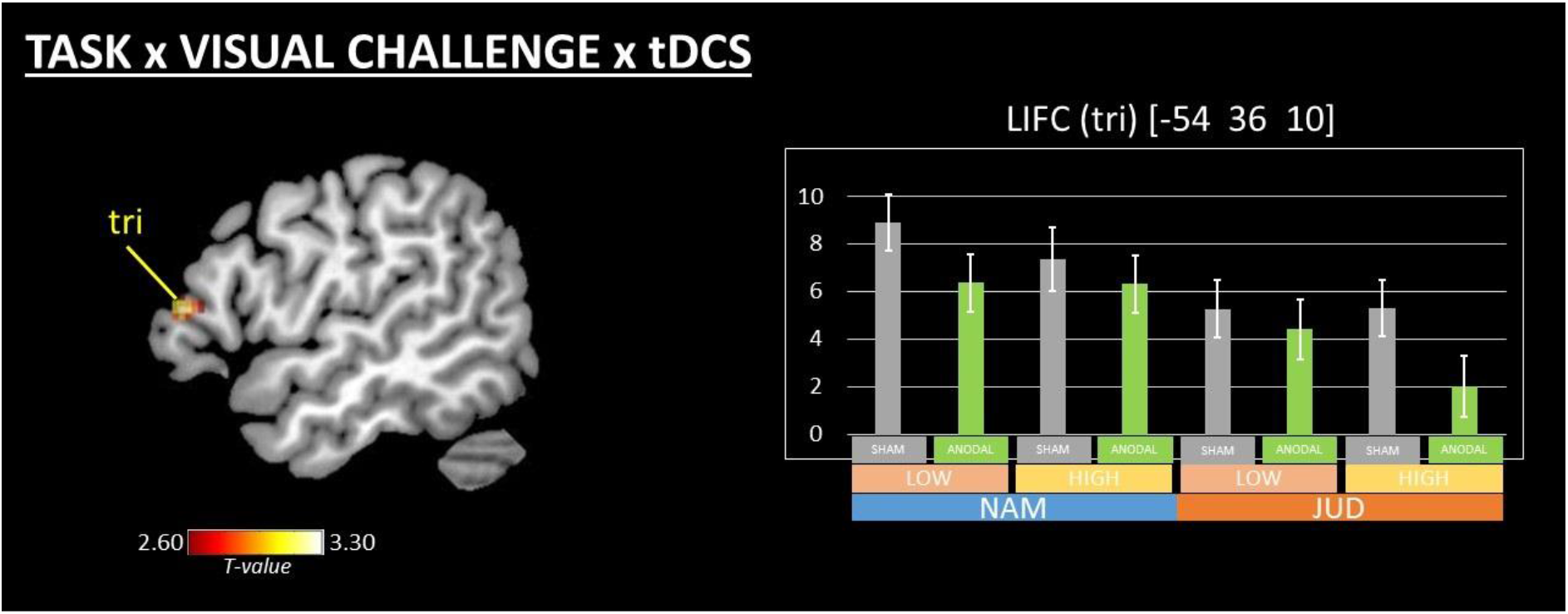
tDCS-related modulations. Areas activated by the Task x Visual challenge x tDCS interaction. Activation is shown at p <0.005-unc. for display purposes. The peak shown survives small-volume correction (cf. main text). Legend: tri = triangular part of the LIFC; NAM = Naming; JUD = Judgment; SHAM = sham tDCS; ANODAL = anodal tDCS; LOW = Low-challenge; HIGH = High-challenge. Y axis in plots shows effect size of BOLD response in arbitrary units.

#### MAIN EFFECTS

A significant main effect of ***tDCS*** (***Anodal>Sham***) was found in the left visual cortex (x y z = −28 −80 6; cluster size = 13; T = 5.25; p = 0.008 FWE-corr.) showing reduced BOLD response during Anodal tDCS with respect to Sham tDCS across tasks and cognitive challenge conditions.

In summary, imaging results showed that: i) the Naming and the Judgment tasks extensively overlapped in the LIFC, although some specific sub-regions within it were more activated by the Naming task; ii) activation patterns in three sub-regions in the LIFC showed that – as compared with Low-challenge conditions – High-challenge conditions were associated with increased BOLD response in the Naming task, but with decreased BOLD response in the Judgment task, in striking contrast with activation patterns in all other MDS nodes (which showed High>Low in both tasks); iii) a neural effect of tDCS was found in the LIFC, further modulated by Task and Visual challenge.

## DISCUSSION

The present study investigated the evidence of domain-specific and domain-general regions within the LIFC, by modulating task domain, cognitive challenge, and Anodal tDCS delivered to the LIFC. The effect of task domain showed that – in the context of matched behavioural performance – a linguistic and a non-linguistic task engaged a common widespread portion of the LIFC, although part of this shared neural substrate was significantly more active in the linguistic task. Behaviourally, the effect of cognitive challenge showed that – as predicted – High-challenge conditions had slower RTs with respect to Low-challenge conditions across both tasks and sensory modalities. Functionally, this was mirrored in all MDS nodes, with the notable exception of the LIFC, where three different sub-regions showed opposite activation patterns. Compared to Low-challenge conditions, High-challenge conditions were associated with an increased BOLD response in the Naming task, but a decreased BOLD response in the Judgment task. Importantly, these results show that these sub-regions within the LIFC have a unique functional profile in the brain, one that suggests domain-specificity, with a preferential processing for Naming. Finally, Anodal tDCS delivered to the LIFC showed a significant modulation of both behaviour and brain, across tasks, and an interaction with Visual challenge. Behaviourally, it reduced RTs for Low-challenge but not High-challenge conditions across both tasks, i.e. what was easy with Sham was made even easier with Anodal tDCS. Functionally, it resulted in reduced BOLD response in a specific sub-region within the LIFC (triangular part) modulated in interaction with Task and Visual challenge.

### Effect of task

Our data show that both a linguistic and a behaviourally-matched non-linguistic task using exactly the same stimuli and type of response recruited a common widespread neural network including several brain areas, and – notably – a large portion of the LIFC (comprising the opercular, triangular, and orbital parts). Such overlapping activation across tasks is consistent with a domain-general view of the functional role of the LIFC. However, it is still possible that within this extensive region, smaller peaks associated with either linguistic or non-linguistic processing are present in slightly different locations across subjects due to inter-subject variability in functional neuroanatomy (cf. Fedorenko et al., 2012, 2013). Our study was not designed to address the issue of inter-subject variability in functional neuroanatomy, but rather to investigate modulatory effects at the group level in order to make inferences about the general population. As such, at the group level overall the LIFC is crucially involved in both tasks.

Nevertheless – notwithstanding the behavioural matching – a portion of the overlapping region in the LIFC (i.e., ventral/posterior opercular and triangular parts) showed higher activation in the Naming than in the Judgment task (cf. ***Naming>Judgment*** contrast; Figure 3B), an evidence of domain-specificity in these sub-regions. In language-specific areas within the LIFC, Fedorenko et al. (2012) have previously reported negative and/or flattened activity patterns associated with domain-general processing (i.e., little-to-no involvement in non-linguistic tasks) across different tasks and stimuli. In the present study, we considered it critical to match task response (overt speech) and stimuli characteristics for across domain comparison (i.e., Y/N vs. Naming). In particular, the use of a cognitive challenge framework (Low-challenge vs. High-challenge) allowed uncertainty in speech response parameters (RTs) to be taken into account when examining effects across domains. This meant that both tasks were likely to rely on a shared circuitry for selecting a spoken response, but within this speech network there would likely be a gradient in which the Naming task would be more likely to rely on language-specific components than the Judgment task. By gradient here we mean an increase in the magnitude of activation observed during the Naming task as opposed to the Judgment task. Therefore, the greater increase in activation during Naming might be related to the increased language selection resources needed to select the target word among a higher number of lexical competitors (Thompson-Schill et al., 1997; Snyder et al., 2007; Rodd et al., 2010a; January et al., 2009; Vitello et al., 2014; Hsu et al., 2017), compared to the Judgement task where the cognitive ambiguity of the decision was varied (RTs), but lexical competition demand was low throughout, i.e. binary Yes/No response.

### Effect of cognitive challenge

In the present study, our main result is that three different sub-regions within the LIFC (located in the opercular, triangular, and orbital parts) exhibit exactly the same modulatory pattern irrespective of the sensory modality involved (i.e., a genuine effect of cognitive challenge, unrelated to the stimulus material at hand). Accordingly, as compared to Low-challenge conditions, in High-challenge conditions activity in these areas increased in the Naming task, whereas it decreased in the Judgment task (cf. Figure 4A-B). Importantly, this was observed in conjunction with RT data that clearly show a consistent effect (i.e., slower responses for High-challenge conditions) in both tasks and sensory modalities (cf. Figure 2F-G), consistent with subjects performing a harder, more demanding task. In both tasks and sensory modalities, High-challenge (i.e., increased cognitive demand) is associated with a higher degree of ambiguity in identifying the various items. However, such an ambiguity has a differential impact on semantic search in the two tasks, namely retrieving the exact linguistic label of a given object in Naming vs. retrieving overall visuo-spatial characteristics in Judgment. Our data suggest that these specific sub-regions in the LIFC are associated with the former (but not the latter) process.

The evidence of opposite cognitive challenge modulatory patterns in the LIFC across the two tasks suggest that these sub-regions are recruited in differing ways. These showed a preference for the Naming task involving linguistic processing (and a sensitivity to cognitive challenge in that specific domain). At the same time, they showed what could be interpreted as the presence of a suppression-like mechanism in the non-linguistic Judgment task. Such an activation pattern has been exhibited by regions relatively disengaged from a specific ongoing task (Merabet et al., 2007; Hairston et al., 2008; Linke et al., 2011; Farooqui & Manly, 2017; Farooqui et al., 2018). Consistent with this and the behavioural data, the LIFC BOLD reduction pattern observed during High-challenge conditions of the Judgment task was accompanied by increased activation elsewhere in the MDS (cf. Figure 5 and see below), as well as in sensory cortices (cf. Supplementary Figure S1C-D). This suggests that further cognitive resources were recruited for the Judgment task in different MDS brain regions as cognitive challenge increased (Duncan, 2010).

Notably, we did not identify any sub-region within the LIFC showing a domain-general modulatory effect. That is, behavioural performance based on cognitive challenge per se, irrespective of task, did not allow us to explain activity in the LIFC. However, it was possible to explain activity based on behavioural performance when considering Naming alone.

In other words, within the LIFC for the Naming task there was consistency between behaviour and BOLD response, whereas in the Judgment task no such consistency was observed. However, within the right inferior frontal cortex (opercular part; cf. Figure 4C and Table 3), we did observe a significant modulation of challenge (High-vs. Low) in the auditory domain, irrespective of task. This suggests different functioning rules in homologue inferior frontal regions across the two hemispheres (e.g., see Cai et al., 2013), the left domain-specific vs. the right domain-general.

This functional discrepancy between the hemispheres may help clarify recovery of left-hemisphere damaged patients with aphasia. In these patients, lesions to the LIFC (or its functional disconnection) are highly likely to impair linguistic processing in domain-specific nodes. Several sources of evidence report that perilesional brain tissue in the LIFC is key for a significant, long-lasting speech recovery (Fridriksson, 2010; Fridriksson et al., 2012). Our present results highlight that the LIFC is not a homogenous functional unit and refine that prediction such that not all perilesional areas within the LIFC might play an equivalent role in speech recovery (i.e., depending on whether perilesional areas include language-specific sub-regions, or not). This might also explain why some patients with more focal left frontal damage are more likely to have domain-specific spoken language deficits (such as anomia) whereas others with more extensive damage show both language and domain-general deficits.

Furthermore, our data suggest that the right inferior frontal cortex may well play a facilitatory role in spoken language production, especially when supported by auditory cues (Blasi et al., 2002; Crinion & Price, 2005; Nardo et al., 2017). Indeed, its domain-general functional characteristics – as indicated in the present study (cf. the contrast ***High-challenge>Low-challenge*** in the Auditory modality) – suggest that this substrate might be sufficiently flexible to support a certain degree of linguistic re-learning (Raboyeau et al., 2008; Richter et al., 2008), although it probably cannot become as efficient as a specialised, hard-wired substrate such as the LIFC (cf. Hartwigsen & Siebner, 2013; Riès et al., 2016).

Our imaging results also showed that – as predicted by the MDS theory (Duncan & Owen, 2000; Duncan, 2010, 2013) – High-challenge (i.e., increased cognitive demand irrespective of task) was associated with increased activation in several MDS nodes (cf. Figure 5 and Supplementary Table ST1), as well as in visual cortices (Supplementary Figure S1C-D and Table 3). Mirroring our behavioural data, in all MDS nodes a greater modulatory effect was observed in the Naming task, (cf. Task x Auditory challenge and Task x Visual challenge, where the difference between High- and Low-challenge is larger in the Naming than in the Judgment task). This shows how sensitive the MDS is to cognitive challenge when processing linguistic material, nicely complementing previous works with non-linguistic material (Fedorenko et al., 2013).

### Effect of tDCS

tDCS has been applied to different brain areas to investigate neuromodulatory effects on various cognitive tasks (Chen et al., 2014; Conson et al., 2015; Pripfl & Lamm, 2015; Brezis et al., 2016; Zmigrod et al., 2016; Payne & Tsakiris, 2017). To our knowledge, the present study is the first to utilise anodal tDCS delivered to the LIFC to directly test its contribution to domain-specific (language) vs. domain-general (cognitively demanding) functioning.

Behaviourally, we found no main effect of anodal tDCS. Instead, we found a significant interaction between Anodal tDCS, cognitive demand (low visual challenge) and task (Naming), with a significant behavioural facilitation, i.e. reduced RTs compared to Sham. That is, if it was easy to Name – during Sham – it was even easier and more efficient (as indexed by faster RTs) when paired with Anodal tDCS delivered to the LIFC. This behavioural difference in tDCS outcomes between two visually different conditions may be related to the novelty of naming our visually challenging (ambiguous) stimuli. Previous research suggests that Anodal tDCS may induce facilitation when the task is well-trained or familiar, but such facilitation is not present during the performance of a novel task (Dockery et al., 2009), or is in accordance with the level of executive control demands (Hussey et al., 2015).

Within the targeted LIFC, a significant neural effect of tDCS was observed for the same interaction with Task and Visual challenge, with a larger decrease in BOLD response for the Naming task during Low-challenge conditions (cf. plot in Figure 6). This effect replicates previous findings from our group (Holland et al., 2011), where concurrent Anodal tDCS paired with a naming task resulted in reduced BOLD response in the same LIFC cluster (triangular part). tDCS itself cannot induce an over-threshold depolarisation of neurons directly, rather it induces firing in neurons that are already near-threshold, which means that neurons unaffected by the task are less likely to discharge (Miniussi & Ruzzoli, 2013). The combination of Anodal tDCS with naming is similar to the co-activation of a specific LIFC network, modulating ongoing long-term potentiation-like changes that outlast the stimulation, leading to consolidation of naming changes and thereby facilitating processing (Miniussi et al., 2013). This is evocative of Hebbian-like plasticity mechanisms. In our (unfamiliar) Judgment task the context is different: the variability of the task to engage the LIFC likely meant variability of the synaptic input function, implying that there was more background noise in the system and little consolidation of the neural networks. In this case, Anodal tDCS would increase both the signal and the noise in the system, both being close to threshold. In this sense, Anodal tDCS delivered to the LIFC would not consistently perturb the neural system supporting the judgment processes. In sum, tDCS requires ongoing learning in order to promote or modify plasticity to prime the task-engaged system and produce corresponding specific effects in the cognitive system, hence the observed interaction effects in the LIFC with task and cognitive challenge.

Notably, only in the neuroimaging data did we find a significant main effect of Anodal tDCS. This was not in the targeted LIFC, but remotely in the left visual cortex, where Anodal tDCS resulted in decreased BOLD response across both tasks irrespective of cognitive demand. Similar remote effects in non-invasive brain stimulation have been reported previously in the motor and language domain (Ward et al., 2010; Antal et al., 2011; Hartwigsen et al., 2017; Fiori et al., 2018). A proposed mechanism of Anodal tDCS is the reduction of the amount of excitatory input required to produce a given response in a task-related (i.e., state-dependent) way (Polania et al., 2018) via modification of synaptic thresholds (i.e., by depolarising neurons close to the firing threshold; see Nitsche & Paulus, 2000, 2001). Increased excitability is associated with reduced BOLD response (i.e., less synaptic input to elicit a given output; cf. Antal et al., 2011; Holland et al., 2011; Fiori et al., 2018). Hence, we interpret this tDCS result in terms of a ‘neural priming’ in the visual cortex. Complex behaviours like naming and making size judgment about objects recruit large-scale bilateral neural systems, and visual processing of the stimuli is the key input to both networks. Therefore, Anodal tDCS is likely to modulate task-related connectivity of regions distant to the stimulation site, as well as task-related areas beneath the electrodes (Boros et al., 2008; Romero Lauro et al., 2014; Vecchio et al., 2018). This implies that the net behavioural effects we observed with Anodal tDCS for both tasks are likely based on a remodelling of the whole task-engaged networks; i.e., complex distributed network interactions rather than being caused by changes of a single left frontal region.

## CONCLUSIONS

Our behavioural, neuroimaging and neuromodulation study indicates a more nuanced picture of domain-specificity vs. domain-generality in the LIFC functioning than previous studies, which have tended to argue for either domain-specific or domain-general aspects of spoken language processing, but not both. Importantly, by factoring out variations in task performance by matching tasks and stimuli characteristics when measuring speech responses (RTs), it allowed meaningful comparisons across domains (e.g., Y/N vs. Naming). In particular, the use of a cognitive challenge framework (easy vs. hard) allowed uncertainty in speech responses to be taken into account when examining response demands across domains. First, our neuroimaging data revealed sub-regions within the LIFC (particularly ventral opercular and triangular parts) that were more strongly activated in a domain-specific manner by the Naming task (language) than the Judgement task. Second, no sub-region within the LIFC was modulated in a domain-general manner by cognitive challenge *per se* (i.e., irrespective of the task at hand). Rather, three different sub-regions within the LIFC (opercular, triangular, and orbital parts) showed a sensitivity to challenge modulation selectively during the Naming task, but not during Judgment. This observed change in magnitude of activation across tasks suggests that there may exist a gradient in which some speech tasks (such as Naming) are more likely to rely on a specific LIFC circuitry for language processing than others (such as making Yes/No decisions). Third, anodal tDCS targeting the LIFC delivered concurrently with both tasks showed a further modulation and consolidation of this domain-specific neural pattern resulting in behavioural changes (RTs).

Taken together, our results highlight the role of specific sub-regions within the LIFC in spoken language production. Within the LIFC we observed a functionally specific neural pattern qualitatively different from all other MDS nodes (with the exception of a small sub-region within the left pre-supplementary motor area; cf. Hertrich et al., 2016), i.e. preferentially related to linguistic processing (Fedorenko et al., 2012, 2013). How the functionally specific sub-network within the LIFC interacts with large-scale MDS (cf. Hsu et al., 2017) remains to be resolved. Future studies might profitably make use of effective connectivity analyses to help clarify the dynamic relationship of the different sub-regions of the LIFC with one another, as well as with the other MDS nodes (Hagoort, 2014; Holland et al., 2016). Observing how these interactions occur may help in identifying how to better support spoken language performance across individuals with language disorders, not just aphasia.

Our results also prepare the ground for possible clinical implications. It has been shown that, following a behavioural anomia treatment, aphasic patients exhibit a significant, robust and long-lasting improvement in speech production that is accompanied by neural priming effects (i.e., reduced BOLD response) in several MDS nodes, including the right homologue of the LIFC (Nardo et al., 2017). Notably, it is not clear whether such a treatment works by improving purely linguistic skills (i.e., domain-specific), or rather general cognitive resources. If adopted with aphasic patients, our protocol – including neurostimulation – might help to disentangle this issue, opening new perspectives to aphasia treatment and outcomes.

## ACKNOWLEDGMENTS

We wish to thank Mr Henry Coley-Fisher and Dr Sheila Kerry for their help with stimuli creation. This study, including JC, DN, KP were funded by a Wellcome Trust Senior Research Fellowship in Clinical Science awarded to JC (106161/Z/14/Z). Scanning was conducted at the Wellcome Trust Centre for Human Neuroimaging (WTCN). The WTCN is supported by core funding from the Wellcome Trust [203147/Z/16/Z].

## SUPPLEMENTARY MATERIAL

### Performance in the Naming task

In the present study, our main behavioural measure was reaction times (RTs). RTs have been computed on all responses provided irrespective of performance. In the Naming task, performance was scored with reference to the target word to-be-named. In case the response provided was different from the target word, the type of response was categorised. Types of response and their occurrence throughout the experiment are reported in Supplementary Table ST2. Overall, we considered four types of response: 1) CORRECT; 2) RELATED; 3) INCORRECT; and 4) MISSING. CORRECT responses included: *target words* (e.g., “cat” for cat), *self-corrections* (“cow… sheep” for sheep), *multiple words* (“apple core” for core), and *phonemes plus target words* (“/sh/… box” for box). RELATED responses included: *super-/sub-ordinate categories* (“bird” for owl), *semantic errors* (“reindeer” for moose), *visual errors* (“snake” for lead), *synonyms* (“present” for gift). INCORRECT responses included: *neologisms, single phonemes without target words* (“/s/” for step), *wrong responses* (“dress” for ball). MISSING responses were those where no response was provided. In the Judgment task, performance could not be assessed in the same way, because there was no ‘target response’ to compare the performance with.

**Supplemental Figure S1.**
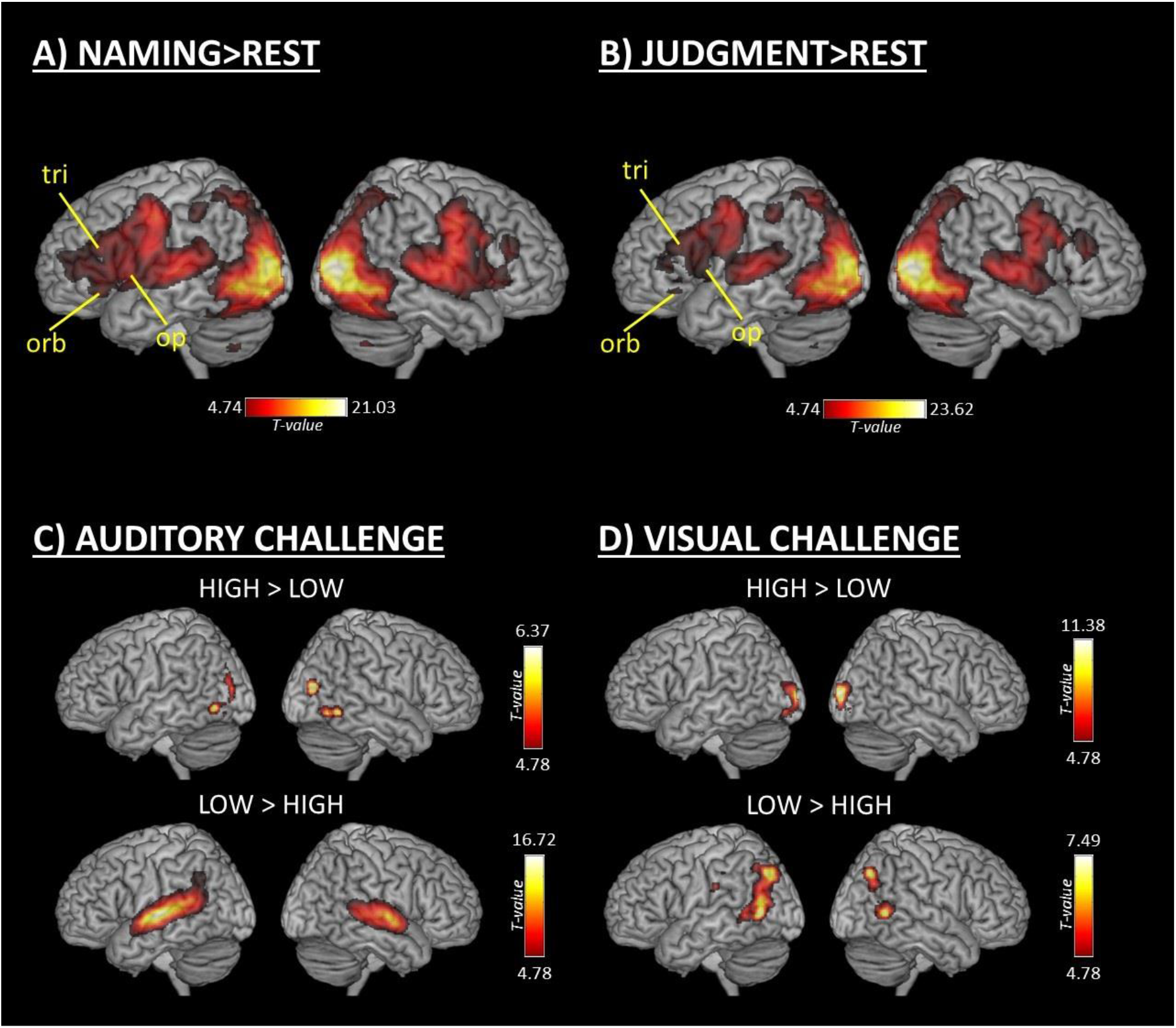
A) Areas associated with performing the Naming task. B) Areas associated with performing the Judgment task. C) Areas associated with Auditory challenge. D) Areas associated with Visual challenge. Legend: op = opercular part of the LIFC; tri = triangular part of the LIFC; orb = orbital part of the LIFC; LOW = Low-challenge; HIGH = High-challenge.

**Supplementary Table ST1 -.**
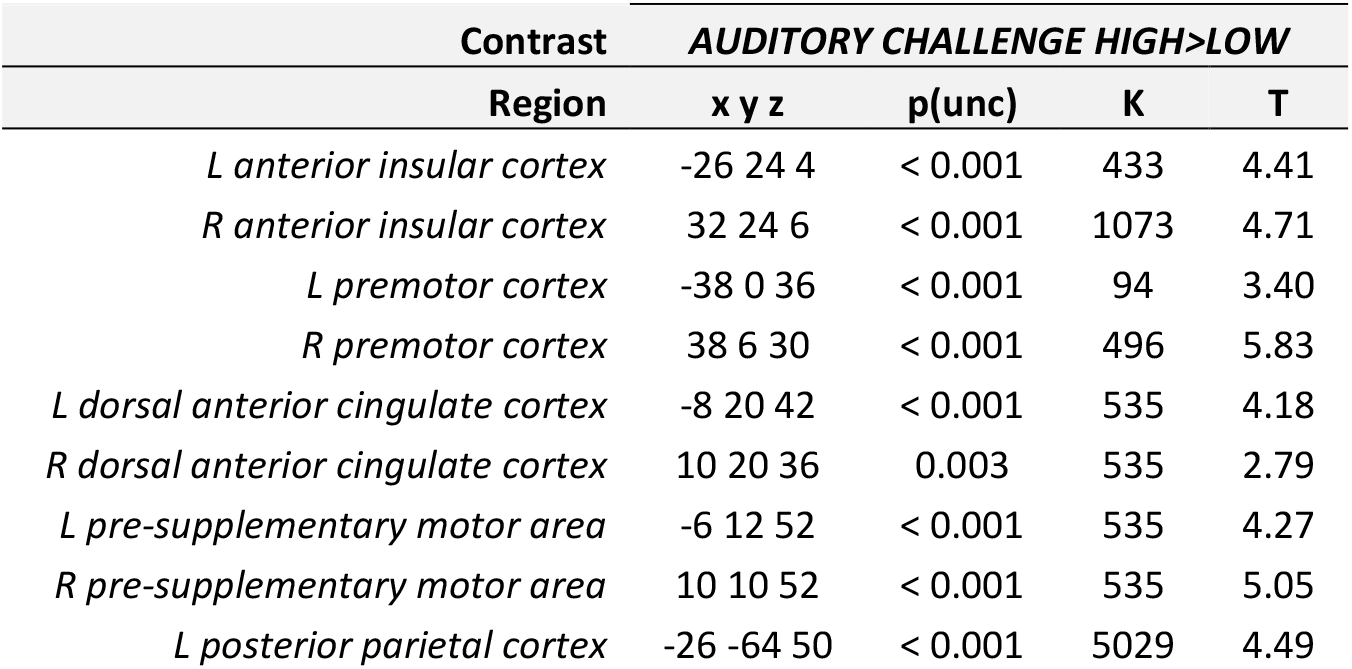
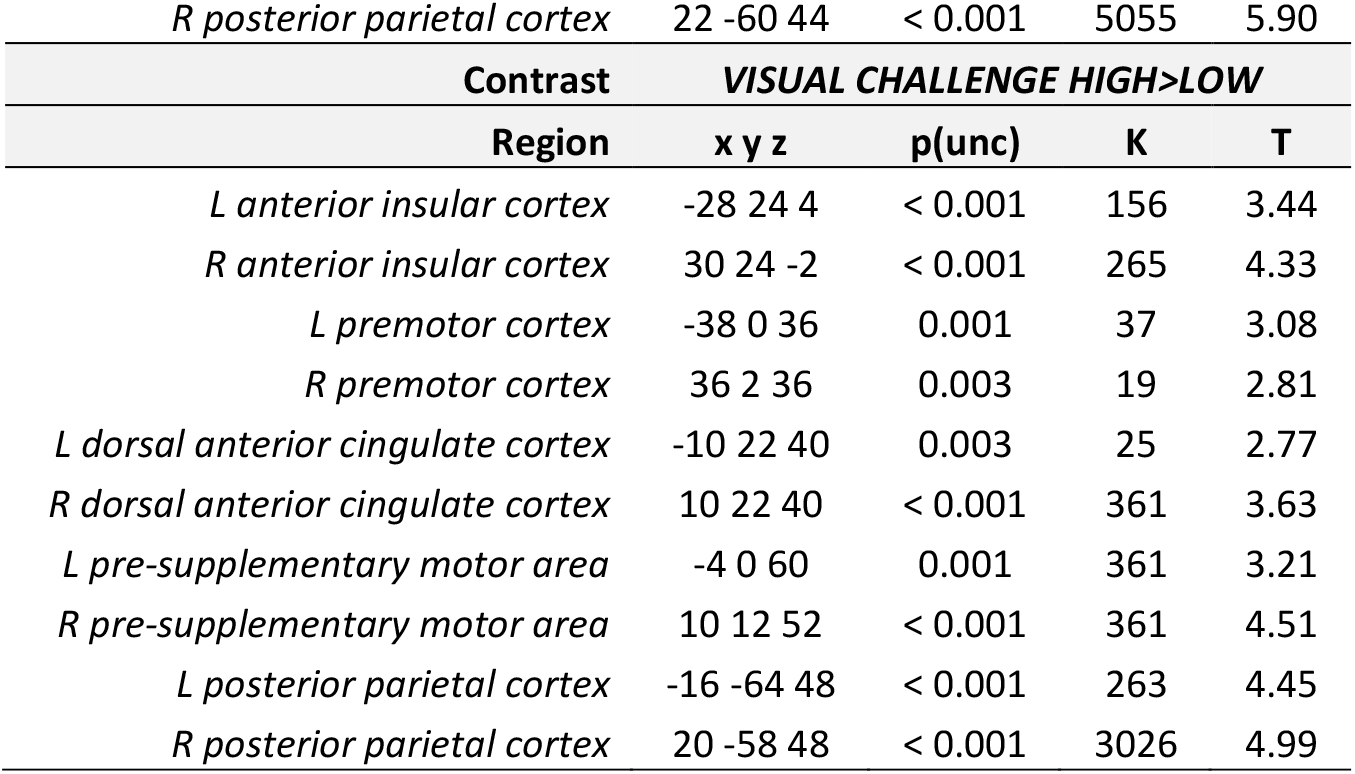
Cognitive challenge-related modulations in other MDS nodes. Legend: R = right; L = left; x y z = MNI coordinates; K = cluster size; T = t-scores; p(unc) uncorrected p-values

**Supplementary Table ST2 -.** Types of response in the Naming task, and their occurrence (in %) throughout the experiment.

| tDCS | SHAM |  |  |  | REAL |  |  |  |  |
| --- | --- | --- | --- | --- | --- | --- | --- | --- | --- |
| VISUAL CHALLENGE | LOW |  | HIGH |  | LOW |  | HIGH |  |  |
| AUDITORY CHALLENGE | LOW | HIGH | LOW | HIGH | LOW | HIGH | LOW | HIGH | mean |
| target words | 86.2 | 56.7 | 75.9 | 45.1 | 84.1 | 58.4 | 67.9 | 39.6 | 64 |
| self-corrections | 0.2 | 1.2 | 0.4 | 0.8 | 0.3 | 1.0 | 0.3 | 0.7 | 1 |
| multiple words | 0.4 | 1.1 | 0.2 | 0.8 | 0.5 | 1.4 | 0.2 | 1.6 | 1 |
| phonemes plus targets | 0.3 | 0.4 | 0.4 | 0.5 | 0.4 | 0.6 | 0.0 | 0.6 | 0 |
| TOT CORRECT | 87.1 | 59.4 | 76.9 | 47.2 | 85.3 | 61.4 | 68.4 | 42.5 | 66 |
| super-/sub-ordinates | 0.1 | 1.4 | 0.4 | 1.4 | 0.4 | 1.1 | 0.1 | 1.1 | 1 |
| semantic errors | 5.0 | 21.1 | 6.2 | 20.6 | 6.1 | 20.3 | 9.2 | 21.9 | 14 |
| visual errors | 3.3 | 8.9 | 3.3 | 12.4 | 2.9 | 9.3 | 5.9 | 13.8 | 7 |
| synonyms | 0.1 | 1.4 | 0.1 | 1.1 | 0.2 | 0.8 | 0.0 | 0.6 | 1 |
| TOT RELATED | 8.5 | 32.8 | 10.0 | 35.5 | 9.6 | 31.5 | 15.2 | 37.4 | 23 |
| neologisms | 0.0 | 0.0 | 0.4 | 0.1 | 0.0 | 0.0 | 0.9 | 0.3 | 0 |
| single phonemes | 0.1 | 0.4 | 0.5 | 0.8 | 0.4 | 0.4 | 1.1 | 0.4 | 1 |
| wrong responses | 3.6 | 5.6 | 9.0 | 11.9 | 3.7 | 5.5 | 10.8 | 15.2 | 8 |
| TOT INCORRECT | 3.7 | 6.0 | 9.9 | 12.8 | 4.1 | 5.9 | 12.8 | 15.9 | 9 |
| MISSING | 0.7 | 1.9 | 3.2 | 4.7 | 1.0 | 1.3 | 3.6 | 4.2 | 3 |
| TOT | 100 | 100 | 100 | 100 | 100 | 100 | 100 | 100 |  |

## REFERENCES

Antal A, Polania R, Schmidt-Samoa C, Dechent P, Paulus W. Transcranial direct current stimulation over the primary motor cortex during fMRI. Neuroimage, 2011;55:590–596.

Bartley JE, Boeving ER, Riedel MC, Bottenhorn KL, Salo T, Eickhoff SB, Brewe E, Sutherland MT, Laird AR. Meta-analytic evidence for a core problem solving network across multiple representational domains. Neurosci Biobehav Rev. 2018;92:318–337.

Blank I, Kanwisher N, Fedorenko E. A functional dissociation between language and multiple-demand systems revealed in patterns of BOLD signal fluctuations. J Neurophysiol. 2014;112(5):1105–18.

Blasi V, Young AC, Tansy AP, Petersen SE, Snyder AZ, Corbetta M. Word retrieval learning modulates right frontal cortex in patients with left frontal damage. Neuron. 2002;36(1):159–70.

Boros K, Poreisz C, Münchau A, Paulus W, Nitsche MA. Premotor transcranial direct current stimulation (tDCS) affects primary motor excitability in humans. Eur J Neurosci. 2008 Mar;27(5):1292–300.

Brezis N, Bronfman ZZ, Jacoby N, Lavidor M, Usher M. Transcranial Direct Current Stimulation over the Parietal Cortex Improves Approximate Numerical Averaging. J Cogn Neurosci. 2016;28(11):1700–1713.

Cai Q, Van der Haegen L, Brysbaert M. Complementary hemispheric specialization for language production and visuospatial attention. Proc Natl Acad Sci U S A. 2013;110(4):E322–30.

Camilleri JA, Müller VI, Fox P, Laird AR, Hoffstaedter F, Kalenscher T, Eickhoff SB. Definition and characterization of an extended multiple-demand network. Neuroimage. 2018;165:138–147.

Chen JC, Hämmerer D, Strigaro G, Liou LM, Tsai CH, Rothwell JC, Edwards MJ. Domain-specific suppression of auditory mismatch negativity with transcranial direct current stimulation. Clin Neurophysiol. 2014;125(3):585–92.

Conson M, Errico D, Mazzarella E, Giordano M, Grossi D, Trojano L. Transcranial Electrical Stimulation over Dorsolateral Prefrontal Cortex Modulates Processing of Social Cognitive and Affective Information. PLoS One. 2015;10(5):e0126448.

Crinion J, Price CJ. Right anterior superior temporal activation predicts auditory sentence comprehension following aphasic stroke. Brain. 2005;128(Pt 12):2858–71.

Dapretto M, Bookheimer SY. Form and content: dissociating syntax and semantics in sentence comprehension. Neuron. 1999 Oct;24(2):427–32.

Darkow R, Martin A, Würtz A, Flöel A, Meinzer M. Transcranial direct current stimulation effects on neural processing in post-stroke aphasia. Hum Brain Mapp. 2017;38(3):1518–1531.

Dockery CA, Hueckel-Weng R, Birbaumer N, Plewnia C. Enhancement of planning ability by transcranial direct current stimulation. J Neurosci. 2009;29(22):7271–7.

Duncan J, Owen AM. Common regions of the human frontal lobe recruited by diverse cognitive demands. Trends Neurosci. 2000 23(10):475–83.

Duncan J. The multiple-demand (MD) system of the primate brain: mental programs for intelligent behaviour. Trends Cogn Sci. 2010 14(4):172–9.

Duncan J. The structure of cognition: attentional episodes in mind and brain. Neuron. 2013 80(1):35–50.

Farooqui AA, Duncan J, Manly T. Hierarchical Cognition causes Task Related Deactivations but not just in Default Mode Regions. bioRxiv 2018 (in press)

Fedorenko E, Thompson-Schill SL. Reworking the language network. Trends Cogn Sci. 2014;18(3):120–6.

Fedorenko E, Hsieh PJ, Nieto-Castañón A, Whitfield-Gabrieli S, Kanwisher N. New method for fMRI investigations of language: defining ROIs functionally in individual subjects. J Neurophysiol. 2010; 104(2):1177–94.

Fedorenko E, Behr MK, Kanwisher N. Functional specificity for high-level linguistic processing in the human brain. Proc Natl Acad Sci U S A. 2011;108(39):16428–33.

Fedorenko E, Duncan J, Kanwisher N. Language-selective and domain-general regions lie side by side within Broca’s area. Curr Biol. 2012 22(21):2059–62.

Fedorenko E, Duncan J, Kanwisher N. Broad domain generality in focal regions of frontal and parietal cortex. Proc Natl Acad Sci U S A. 2013;110(41):16616–21.

Fiori V, Kunz L, Kuhnke P, Marangolo P, Hartwigsen G. Transcranial direct current stimulation (tDCS) facilitates verb learning by altering effective connectivity in the healthy brain. Neuroimage. 2018;181:550–559.

Fridriksson J. Preservation and modulation of specific left hemisphere regions is vital for treated recovery from anomia in stroke. J Neurosci. 2010;30(35):11558–64.

Fridriksson J, Richardson JD, Fillmore P, Cai B. Left hemisphere plasticity and aphasia recovery. Neuroimage. 2012 Apr 2;60(2):854–63.

Geranmayeh F, Wise RJ, Mehta A, Leech R. Overlapping networks engaged during spoken language production and its cognitive control. J Neurosci. 2014;34(26):8728–40.

Hagoort P. Nodes and networks in the neural architecture for language: Broca’s region and beyond. Curr Opin Neurobiol. 2014;28:136–41.

Hartwigsen G, Bzdok D, Klein M, Wawrzyniak M, Stockert A, Wrede K, Classen J, Saur D. Rapid short-term reorganization in the language network. Elife 2017;6:e25964.

Hartwigsen G, Siebner HR. Novel methods to study aphasia recovery after stroke. Front Neurol Neurosci. 2013;32:101–11.

Hertrich I, Dietrich S, Ackermann H. The role of the supplementary motor area for speech and language processing. Neurosci Biobehav Rev. 2016;68:602–610.

Holland R, Leff AP, Josephs O, Galea JM, Desikan M, Price CJ, Rothwell JC, Crinion J. Speech facilitation by left inferior frontal cortex stimulation. Curr Biol. 2011;21(16):1403–7.

Holland R, Leff AP, Penny WD, Rothwell JC, Crinion J. Modulation of frontal effective connectivity during speech. Neuroimage. 2016;140:126–33.

Hsu NS, Jaeggi SM, Novick JM. A common neural hub resolves syntactic and non-syntactic conflict through cooperation with task-specific networks. Brain Lang. 2017;166:63–77.

January D, Trueswell JC, Thompson-Schill SL. Co-localization of stroop and syntactic ambiguity resolution in Broca’s area: implications for the neural basis of sentence processing. J Cogn Neurosci. 2009;21(12):2434–44.

Miniussi C, Ruzzoli M. Transcranial stimulation and cognition. Handb Clin Neurol. 2013;116:739–50.

Miniussi C, Harris JA, Ruzzoli M. Modelling non-invasive brain stimulation in cognitive neuroscience. Neurosci Biobehav Rev. 2013;37(8):1702–12.

Nardo D, Holland R, Leff AP, Price CJ, Crinion JT. Less is more: neural mechanisms underlying anomia treatment in chronic aphasic patients. Brain. 2017 140(11):3039–3054.

Nitsche MA, Paulus W. Excitability changes induced in the human motor cortex by weak transcranial direct current stimulation. J. Physiol 2000;527:633–639.

Nitsche MA, Paulus W. Sustained excitability elevations induced by transcranial DC motor cortex stimulation in humans. Neurology, 2001;57(10):1899–1901.

Noppeney U, Phillips J, Price C. The neural areas that control the retrieval and selection of semantics. Neuropsychologia. 2004;42(9):1269–80.

Novick JM, Kan IP, Trueswell JC, Thompson-Schill SL. A case for conflict across multiple domains: memory and language impairments following damage to ventrolateral prefrontal cortex. Cogn Neuropsychol. 2009;26(6):527–67.

Payne S, Tsakiris M. Anodal transcranial direct current stimulation of right temporoparietal area inhibits self-recognition. Cogn Affect Behav Neurosci. 2017;17(1):1–8.

Petkov CI, Marslen-Wilson WD. Editorial overview: The evolution of language as a neurobiological system. Curr Opin Behav Sci. 2018;21:v–xii.

Polanía R, Nitsche MA, Ruff CC. Studying and modifying brain function with non-invasive brain stimulation. Nat Neurosci. 2018;21(2):174–187.

Price CJ. A review and synthesis of the first 20 years of PET and fMRI studies of heard speech, spoken language and reading. Neuroimage. 2012 62(2):816–47.

Pripfl J, Lamm C. Focused transcranial direct current stimulation (tDCS) over the dorsolateral prefrontal cortex modulates specific domains of self-regulation. Neurosci Res. 2015 Feb;91:41–7.

Raboyeau G, De Boissezon X, Marie N, Balduyck S, Puel M, Bézy C, Démonet JF, Cardebat D. Right hemisphere activation in recovery from aphasia: lesion effect or function recruitment? Neurology. 2008 Jan 22;70(4):290–8.

Richter M, Miltner WH, Straube T. Association between therapy outcome and right-hemispheric activation in chronic aphasia. Brain. 2008;131(Pt 5):1391–401.

Riès SK, Dronkers NF, Knight RT. Choosing words: left hemisphere, right hemisphere, or both? Perspective on the lateralization of word retrieval. Ann N Y Acad Sci. 2016;1369(1):111–31.

Rodd JM, Johnsrude IS, Davis MH. The role of domain-general frontal systems in language comprehension: evidence from dual-task interference and semantic ambiguity. Brain Lang. 2010;115(3):182–8.

Rodd JM, Vitello S, Woollams AM, Adank P. Localising semantic and syntactic processing in spoken and written language comprehension: an Activation Likelihood Estimation meta-analysis. Brain Lang. 2015;141:89–102.

Romero Lauro LJ, Rosanova M, Mattavelli G, Convento S, Pisoni A, Opitz A, Bolognini N, Vallar G. TDCS increases cortical excitability: direct evidence from TMS-EEG. Cortex. 2014;58:99–111.

Santi A, Grodzinsky Y. Working memory and syntax interact in Broca’s area. Neuroimage. 2007;37(1):8–17.

Snyder HR, Feigenson K, Thompson-Schill SL. Prefrontal cortical response to conflict during semantic and phonological tasks. J Cogn Neurosci. 2007;19(5):761–75.

Thompson-Schill SL, D’Esposito M, Aguirre GK, Farah MJ. Role of left inferior prefrontal cortex in retrieval of semantic knowledge: a reevaluation. Proc Natl Acad Sci U S A. 1997;94(26):14792–7.

Tyler LK, Marslen-Wilson WD, Randall B, Wright P, Devereux BJ, Zhuang J, Papoutsi M, Stamatakis EA. Left inferior frontal cortex and syntax: function, structure and behaviour in patients with left hemisphere damage. Brain. 2011;134(Pt 2):415–31.

Tzourio-Mazoyer N, Landeau B, Papathanassiou D, Crivello F, Etard O, Delcroix N, Mazoyer B, Joliot M. Automated anatomical labeling of activations in SPM using a macroscopic anatomical parcellation of the MNI MRI single-subject brain. Neuroimage. 2002;15(1):273–89.

Vecchio F, Di Iorio R, Miraglia F, Granata G, Romanello R, Bramanti P, Rossini PM. Transcranial direct current stimulation generates a transient increase of small-world in brain connectivity: an EEG graph theoretical analysis. Exp Brain Res. 2018;236(4):1117–1127.

Vitello S, Warren JE, Devlin JT, Rodd JM. Roles of frontal and temporal regions in reinterpreting semantically ambiguous sentences. Front Hum Neurosci. 2014;8:530.

Ward NS, Bestmann S, Hartwigsen G, Weiss MM, Christensen LOD, Frackowiak RSJ, Rothwell JC, Siebner HR. Low-Frequency Transcranial Magnetic Stimulation over Left Dorsal Premotor Cortex Improves the Dynamic Control of Visuospatially Cued Actions. J Neurosci 2010;30:9216–9223.

Woolgar A, Bor D, Duncan J. Global increase in task-related fronto-parietal activity after focal frontal lobe lesion. J Cogn Neurosci. 2013;25(9):1542–52.

Zmigrod S, Zmigrod L, Hommel B. Transcranial direct current stimulation (tDCS) over the right dorsolateral prefrontal cortex affects stimulus conflict but not response conflict. Neuroscience. 2016;322:320–5.

